# Flow-Triggered Regeneration of Electrochemical Aptasensor for Continuous Monitoring of 17β-Estradiol

**DOI:** 10.64898/2026.10.05.755148

**Authors:** Hadi Mirzajani, Giacomo Massini, Zahra Davari, Francesca Rodino, Sandro Carrara

## Abstract

Continuous biomarker monitoring with affinity biosensors remains limited by probe saturation, slow dissociation, signal carryover, and regeneration methods that require chemical washing, harsh buffers, or external reset modules. Here, we report a flow-programmed, self-regenerative electrochemical aptamer-based biosensor for continuous 17β-estradiol monitoring. The platform restores the sensing interface by tuning buffer flow through the device, enabling repeated sensing–regeneration cycles without chemical stripping or active-reset hardware. The biosensor achieves a 1.46 pM limit of detection, and a log-linear dynamic range from 100 pM to 1 µM in real human sweat, covering physiologically relevant estradiol concentrations. It exhibits high selectivity, with responses to potential molecularly similar interferents remaining below 10% of the 17β-estradiol signal. Under dynamic flow operation, the device shows reversible signal recovery and stable electrochemical performance, with higher buffer-wash flow reducing baseline carryover between measurements. A coupled binding–transport simulation captures the flow-dependent regeneration trends and maps how wash-flow rate, wash duration, and apparent aptamer affinity affect regeneration efficiency. Ex vivo testing in real human sweat, including samples from pregnant women, supports translational relevance. This work establishes buffer-flow programming as a reagent-free route to regenerable electrochemical aptamer sensing for automated, wearable-compatible estradiol monitoring.

## 1. Introduction

Continuous biosensing is increasingly recognized as essential for capturing physiology as it evolves over time rather than through isolated single-point measurements. For affinity-based sensors, however, the ability to operate continuously is inseparable from the ability to regenerate. Once recognition sites become occupied, the sensing interface must return rapidly and reproducibly to its baseline state; otherwise, measurements become vulnerable to saturation, hysteresis, slow washout, drift, and repeated sensor replacement. This challenge is especially severe for low-abundance analytes, for which high-affinity receptors are often required^1, 2^. Although stronger binding improves sensitivity, it can also impose long complexation half-lives and slow re-equilibration when analyte concentrations decline^2^. Regeneration is therefore not a secondary operational feature, but a central design requirement for any biosensor intended for repeated, longitudinal, or continuous deployment^2–6^.

Aptamer-based electrochemical biosensors are particularly well suited to address this challenge^7, 8^. Since the seminal demonstration of label-free electronic detection using an aptamer-modified electrode^9^, structure-switching electrochemical aptamer-based sensors have developed into a versatile class of reagentless devices in which target binding alters the conformation or flexibility of a redox-labeled nucleic acid layer and thereby modulates electron transfer at the electrode surface^10^. Their direct electrical readout, compatibility with miniaturized electrodes, and ability to function in complex samples make them highly attractive for wearable, implantable, and continuous sensing applications^5, 8, 10^. At the same time, the very affinity interactions that enable sensitive detection can also hinder rapid return to baseline after target capture, making regeneration a central requirement in the design of continuous aptamer sensors^2, 9^.

A number of regeneration strategies have been explored across aptamer biosensor formats. An on-chip structure-switching aptamer platform was demonstrated using ferrocene-labelled IFN-γ aptamers immobilized on magnetic nanobeads and confined within a microfluidic device, enabling continuous electrochemical monitoring in cell culture medium and serum without exogenous reagents^11^. The RAPTER robotic electrochemical aptamer platform was developed for luteinising hormone, showing rapid, repeatable hormone profiling in clinical serum and regeneration to >93% below saturation concentrations^12^. A real-time optical endotoxin aptasensor was reported with ensemble and single-molecule readout and was regenerated for repeated use^13^. Aptamer-based recognition was further translated into a wearable FET smartwatch for real-time sweat cortisol monitoring, supporting aptamer platforms for continuous non-invasive hormone sensing, although regeneration was not the central focus of the study^14^. More recently, active-reset protein sensors have shown that externally applied high-frequency oscillations can accelerate dissociation and regenerate the unbound state^2^. Collectively, these studies establish that regeneration is feasible, but many existing strategies depend on dedicated regeneration buffers, chemically harsh washing steps, or added actuation modules that may complicate integration into simple, wearable, and continuously operating devices^15^.

17β-Estradiol is a particularly important target for such platforms because its dynamic variation is central to female reproductive physiology and hormone monitoring, while current assessment still relies on episodic sampling. Aptamer-based estradiol sensors have evolved from early electrochemical detection on gold electrode chips^16^ to nanocomposite electrochemical aptasensors with improved sensitivity^17^ and label-free impedance biosensors validated in human urine^18^. Importantly, a recent wearable aptamer nanobiosensor demonstrated non-invasive estradiol monitoring in sweat and reported cyclical sweat estradiol fluctuations together with a strong correlation between sweat and blood measurements, underscoring the promise of sweat as a clinically relevant matrix for hormone tracking^19^. What remains needed, however, is a regeneration strategy that is intrinsically compatible with continuous electrochemical operation in sweat rather than appended to the sensor as a separate chemical or mechanical reset step^16^.

In this work, we address this challenge through a new flow-triggered self-regenerative electrochemical aptamer biosensor for continuous monitoring of 17β-estradiol. Rather than relying on external reset hardware or specialized regeneration reagents, our platform regenerates the sensing interface simply by tuning the flow of buffer through the sensor. This approach couples molecular recognition and interface recovery to a single fluidic control parameter, simplifying operation while enabling repeated sensing-regeneration cycles and sustained use. We further demonstrate the translational relevance of this concept through ex vivo validation of biosensor in real human sweat, including sweat samples collected from pregnant women. To support the experimental interpretation, we also develop a coupled binding–transport simulation that incorporates device geometry, chamber exchange, interfacial analyte depletion, surface aptamer occupancy, and current response. The model captures the observed flow-dependent regeneration trends and maps how wash-flow rate, wash duration, and apparent aptamer affinity influence regeneration efficiency. By combining experimental flow-programmed regeneration with simulation-guided design-space analysis, this work establishes a practical framework for reagent-free, wearable-compatible, and semi-continuous estradiol monitoring that can be integrated into future wearable platforms.

## 2. Results

### 2.1. In vitro EIS characterization of aptamer-based biosensor

An E2-selective aptamer-based electrochemical biosensor was first evaluated in a label-free impedimetric format to verify target recognition at the functionalized Au interface. Electrochemical impedance spectroscopy (EIS) was performed in a ferri/ferrocyanide redox system using a surface-immobilized 17β-estradiol-selective aptamer without redox labels, following the established principle that E2 binding to surface-confined aptamers modulates interfacial charge-transfer resistance, *R*_ct_ ^18^. Details of the sensing setup, electrode cleaning, and functionalization characterization are provided in Methods section of SI and Fig. S1. As schematically illustrated in **Figure 1a**, E2 binding induces an aptamer–target complex formation and structural rearrangement of the surface-confined probe layer, thereby altering redox-probe accessibility and charge-transfer behavior at the Au/electrolyte interface^20, 21^. The resulting *R*_ct_ modulation provides a label-free electrochemical readout of E2 recognition and establishes the mechanistic basis for subsequent calibration.

**Figure 1.**
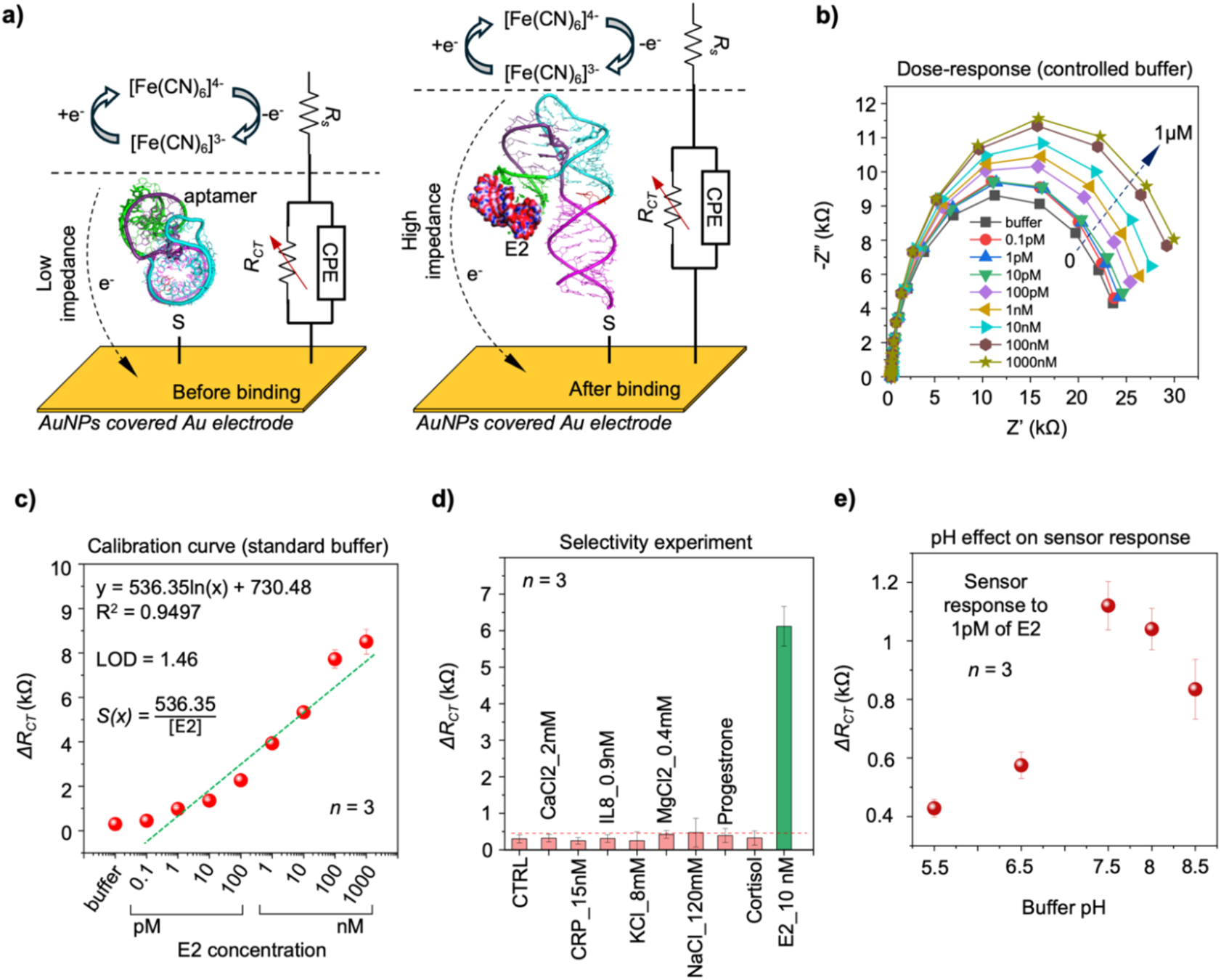
In vitro characterization of the developed biosensor in a controlled buffer. (a) Schematic illustration of sensor operation and aptamer switching in the presence of the target. (b) Dose–response of the sensor to different concentrations of E2 in the controlled buffer. (c) Calibration curve and corresponding calibration equation of the developed biosensor. (d) Selectivity results for eight potential interfering molecules spiked into the buffer. (e) Effect of pH variations of the buffer on the response of the sensor to a 1 pM of estradiol. Each condition was measured at least three times, error bars indicate the standard deviation.

#### 2.1.1. Dose-response experiment and calibration curve extraction in vitro

The working electrode consisted of an Au layer decorated with 40 nm AuNPs, which increased the electroactive surface area and improved interfacial signal transduction, as confirmed by the comparative electrode study in Figure S2 and consistent with the established use of AuNPs for enhancing electrochemical biosensor performance^22^. Concentration-dependent EIS measurements were then performed after incubation with increasing E2 concentrations, and the resulting Nyquist plots are shown in Figure 1b. Measurements were conducted in binding buffer containing 2 mM Tris–HCl, pH 7.5, 10 mM NaCl, 0.5 mM KCl, 0.2 mM MgCl₂, 0.1 mM CaCl₂, and 5% ethanol. Across the tested concentration range of 0.1 pM to 1 µM, the impedance spectra showed a systematic E2-dependent change in interfacial charge-transfer behavior, indicating that target binding to the surface-confined aptamer produced a reproducible modulation of the electrode/electrolyte interface. The extracted calibration curve, obtained by fitting *R*_ct_ values using a standard Randles equivalent circuit model, exhibited a log-linear response from 1 pM to 1 µM, with good linearity (*R*² = 0.9497) and a calculated LOD of 1.46 pM using the 3.3σ/S criterion^23–25^ where σ is the standard deviation of sensor response, and *S* is the slope of the calibration curve. These results demonstrate that the aptamer-functionalized AuNPs-decorated Au electrode enables low-picomolar E2 detection and provides a broad analytical window extending from the sweat-relevant picomolar regime to high challenge concentrations, supporting its use for subsequent ex vivo analysis of pregnant and non-pregnant sweat samples^19^. Furthermore, benchmarking against a commercial ELISA assay using matched E2 solutions showed near-unity agreement within the overlapping concentration range (Figure S3), supporting the analytical reliability of the EIS-based readout.

#### 2.1.2. Selectivity analysis

Selectivity was evaluated by testing the aptamer-functionalized sensor with sweat-relevant ionic species and biomolecular interferents, including CaCl₂ 2 mM, KCl 8 mM, MgCl₂ 0.4 mM, NaCl 120 mM, CRP 15 nM, and IL-8 0.9 nM, as well as structurally related steroid hormones, including progesterone and cortisol, tested at a tenfold higher concentration than E2. As summarized in Figure 1d, compared with the response to 10 nM E2, the non-target steroids produced only minor changes in *ΔR*_ct_, indicating that the observed signal is dominated by E2-specific aptamer recognition rather than nonspecific steroid adsorption or general hydrophobic interference at the electrode surface. The pH dependence of the response was also evaluated to define a stable operating condition, because buffer pH can influence nucleic-acid conformation, aptamer–target binding, and electrochemical signal transduction in aptamer-based sensors^3, 20,21^. As shown in Figure 1e, pH 7.5 provided the most suitable response profile and was therefore selected for subsequent measurements. Overall, these experiments demonstrate that the aptamer recognition layer selectively distinguishes E2 from closely related steroids and relevant interferents, while pH optimization defines the buffer environment required for reliable aptamer operation.

#### 2.1.3. Sensor analysis under sweat-like ionic stress

Because sweat electrolyte composition varies between individuals and within the same individual as a function of sweat rate, physiological state, exercise conditions, hydration, diet, and sampling context, sensor performance was evaluated under elevated sweat-relevant ionic stress conditions^26, 27^. For this analysis, the EIS assay was repeated in separately prepared binding buffers containing high concentrations of selected sweat-relevant ions, including 226 mM NaCl, 5.75 mM CaCl₂, or 9.4 mM MgCl₂. Each buffer was spiked with E2 concentrations ranging from 0.1 pM to 100 nM and tested using the AuNPs-decorated, aptamer-functionalized Au electrodes under the same experimental procedure described for the dose-response calibration in Section 2.1.1. As summarized in **Figure 2a–c**, the aptamer-functionalized interface retained a clear concentration-dependent response to E2 under all ionic conditions, demonstrating that the E2 recognition layer remained operational under simplified high-ionic-strength sweat-like media. However, the calibration parameters were strongly dependent on ion composition. In 226 mM NaCl, the sensor showed strong log-linear behavior (*R*² = 0.9832) with an LOD of 1.21 pM. In 5.75 mM CaCl₂, the response remained linear (*R*² = 0.9497) with an LOD of 0.346 pM, although the reduced slope indicated partial signal damping. In 9.4 mM MgCl₂, the sensor exhibited a larger response slope and low LOD of 0.302 pM, but reduced linearity (*R*² = 0.9171), suggesting stronger ion-dependent variability. These results show that the sensing interface retains E2 responsiveness under elevated Na⁺-, Ca²⁺-, and Mg²⁺-rich conditions, supporting functional tolerance of the biorecognition layer to simplified sweat-like ionic stress. At the same time, the ion-dependent shifts in calibration slope, LOD, and linearity demonstrate that the response is matrix-sensitive rather than matrix-independent. The NaCl-dependent calibration shift is consistent with increased ionic strength compressing the diffuse double layer and enhancing electrostatic/Debye screening at nucleic-acid-functionalized sensing interfaces^28^, whereas the Mg²⁺- and Ca²⁺-dependent responses are consistent with reports that divalent cations can modulate aptamer folding, target recognition, and local charge redistribution in structure-switching aptamer sensors^29, 30^. Importantly, these ionic effects did not abolish E2 recognition; rather, they redistributed the calibration landscape, indicating that quantitative in situ sweat analysis should incorporate proper system-integration with the capability of matrix-matched calibration or ionic-strength correction before real-sample deployment.

**Figure 2.**
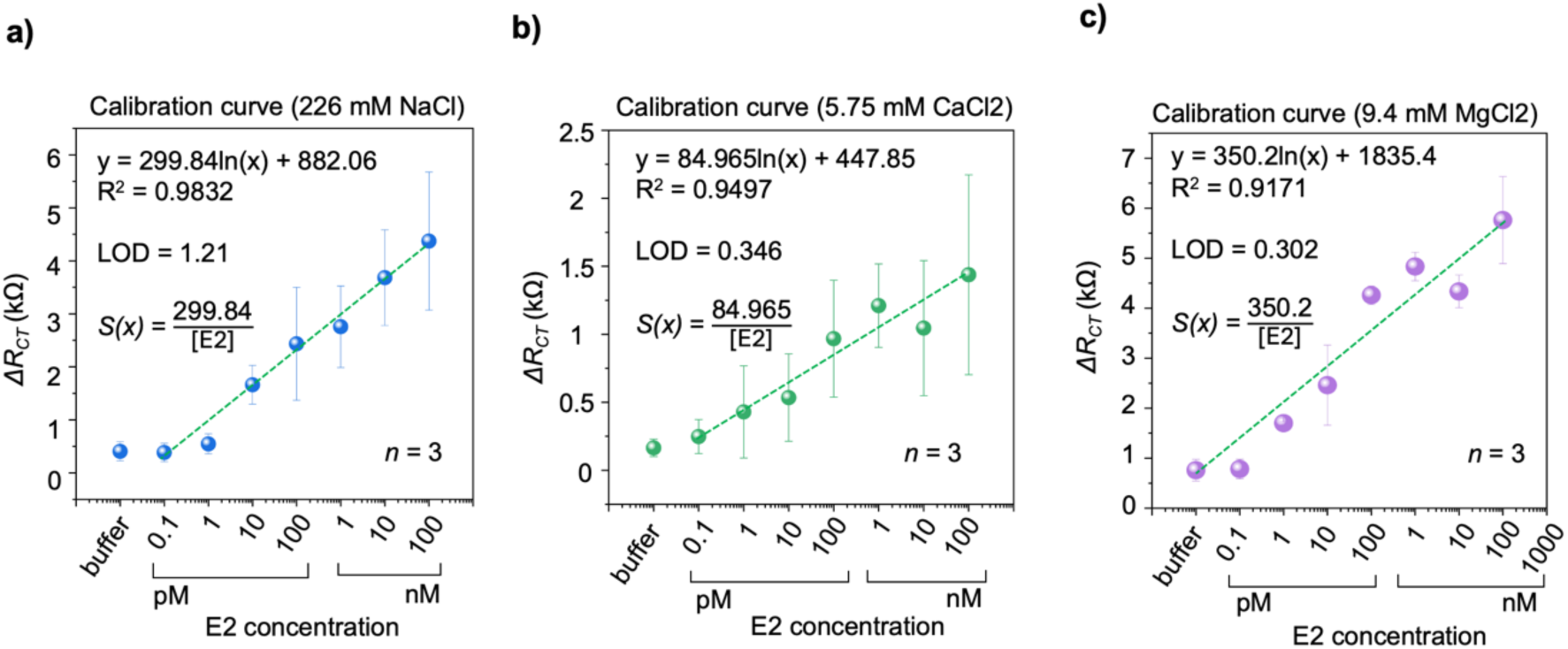
Effect of elevated ionic strength of the binding buffer (2 mM Tris–HCl, pH 7.5, containing 10 mM NaCl, 0.5 mM KCl, 0.2 mM MgCl₂, 0.1 mM CaCl₂, and 5% ethanol) on the in vitro biosensor response. (a) Calibration response in buffer containing 226 mM NaCl, showing strong log-linear behavior. (b) Calibration response in buffer containing 5.75 mM CaCl₂, showing retained E2 responsiveness with reduced signal slope. (c) Calibration response in buffer containing 9.4 mM MgCl₂, showing enhanced response magnitude but greater ion-dependent variability.

#### 2.1.4. Stability and shelf-life characterization Stability characterization

For the stability study, independently functionalized sensors were stored at room temperature (25 °C), refrigerator (4 °C), or elevated temperature (37 °C), all in wet condition (PBS), in sealed containers, and protected from light. At predefined time points, sensors were removed from storage and incubated with 1 nM of E2 in binding buffer and then characterized by EIS in ferri/ferrocyanide redox solution, with *ΔR_ct_* used as the output metric, and the results are plotted in **Figure 3a,b,c**. The shaded ±15% region was used as a practical stability window, corresponding to retention of at least 85% of the initial sensor response and aligned with common analytical acceptance criteria and previously reported biosensor storage-stability benchmarks^31^. Sensors stored at room temperature showed substantial performance loss (Figure 3a). The response began to decrease within the first days and dropped by approximately 57% after 30 days, moving well outside the ±15% stability window. This deterioration can be considered as the result of progressive destabilization of the aptamer-modified gold interface, including possible desorption or reorganization of the thiolated aptamer/blocking monolayer and chemical alteration of the alkanethiolate surface layer. This interpretation is supported by prior electrochemical aptamer-based sensor studies showing storage-condition-dependent aptamer retention and signal loss at room temperature^32^, mechanistic reports demonstrating temperature-dependent dissociation of thiol-modified DNA from gold^33^, and surface-science studies showing ambient oxidative degradation of alkanethiolate self-assembled monolayers on gold^34^. In addition, studies on EAB sensors have shown that the chemistry and terminal groups of alkanethiol monolayers strongly influence long-term signal stability, baseline drift, and sensor response in complex media^35, 36^. These results indicate that ambient storage is unsuitable for long-term preservation. In contrast, refrigerated storage at 4 °C effectively preserved sensor function, maintaining the *ΔR*_ct_ response close to its initial value and within the predefined ±15% stability window for at least 10 days, corresponding to ≥85% signal retention (Figure 3b). This behavior is consistent with improved retention reported for electrochemical aptamer-based sensors under low-temperature storage conditions^32^ and with aptasensor studies in which storage at 4 °C preserved most of the initial electrochemical response over time^37, 38^. Therefore, refrigeration appears to slow early interfacial drift and helps maintain the functional aptamer/blocking layer during short-term storage. Sensors stored at 37 °C retained measurable activity initially, but the response progressively declined and sensor-to-sensor variability increased over time (Figure 3c), suggesting thermally accelerated loss of response stability and reduced analytical reliability. This behavior is consistent with the known temperature sensitivity of thiolated DNA/gold interfaces^33^ and with electrochemical aptamer-based sensor studies identifying monolayer instability, and surface-chemistry-dependent drift as key contributors to signal loss at physiological or elevated temperatures^35, 39^. Together, these results establish a clear temperature dependence in sensor preservation, where refrigeration limits early performance drift and maintains the functional sensing layer, whereas ambient and elevated-temperature storage accelerate signal decay and reduce long-term analytical reliability.

**Figure 3.**
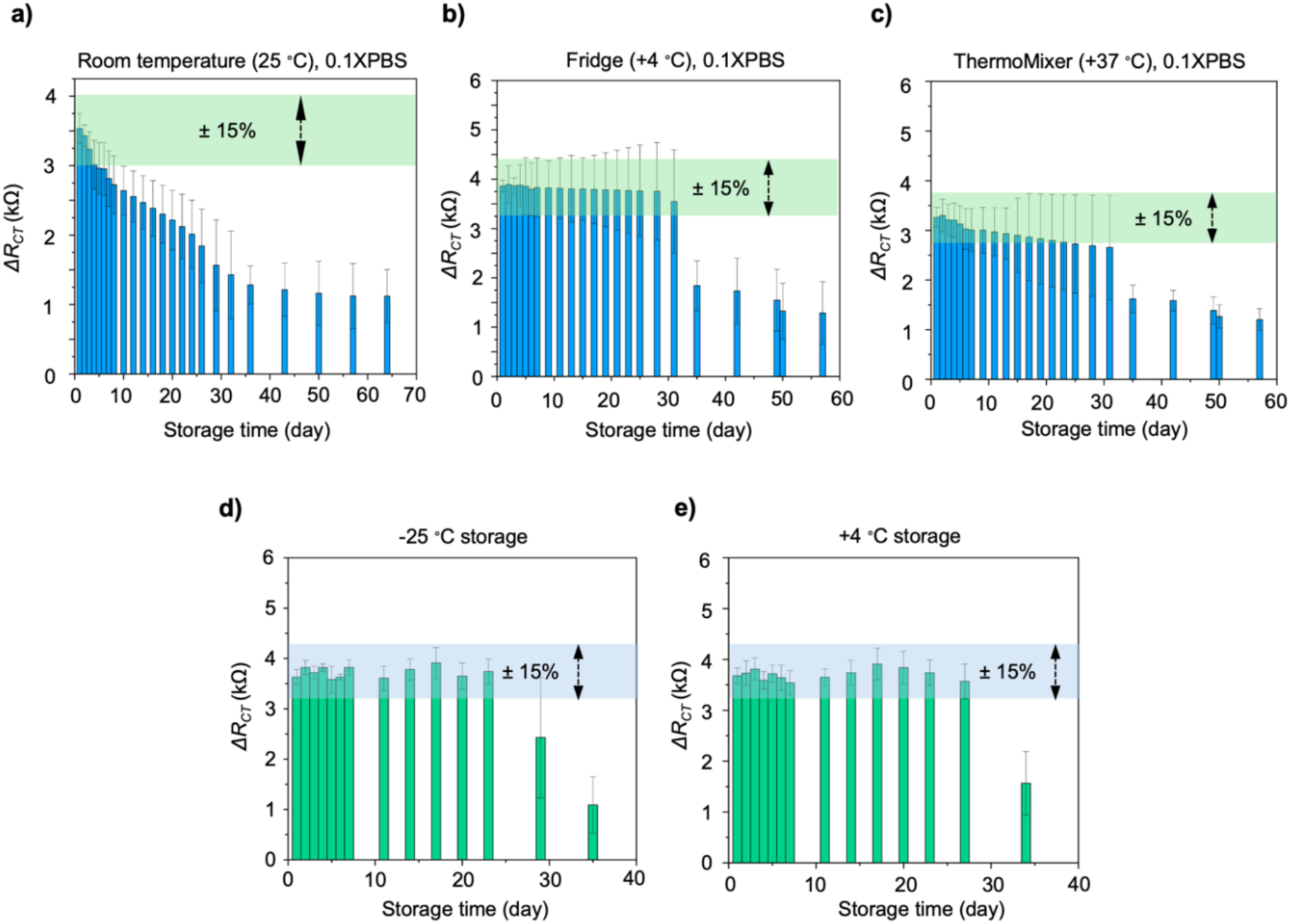
Stability and shelf-life characterization of the biosensor under different storage conditions. (a) Stability during storage at room temperature (25 °C) in 0.1× PBS. (b) Stability during storage at 4 °C in 0.1× PBS. (c) Stability during storage at 37 °C (Thermomixer) in 0.1× PBS. (d) Shelf-life during storage at −25 °C (freezer). (e) Shelf-life during storage at 4 °C (refrigerator). For each condition *n* = 3, and error bars indicate standard deviation.

#### Shelf-life characterization

A shelf-life study was also performed using a larger batch of 60 sensors fabricated and functionalized on the same day. After functionalization, sensors were stored at either −25 °C or 4 °C and periodically tested using the same EIS protocol. As shown in Figure 3d,e, both cold-storage conditions preserved sensor performance within the ±15% window for approximately 30 days, with comparable average response and variability. Experiments indicate around two days more stability for sensors stored in 4 °C. However, after longer storage (around 30 days), the response decreased sharply for both storage conditions, indicating eventual loss of activity. This study indicates that low-temperature storage improves long-term retention of sensor function, including aptamer retention, signal gain, and affinity, and with reports identifying degradation of thiolated aptamer monolayers on gold as a key factor limiting long-term electrochemical aptamer-sensor stability^32, 39^. It is also supported by earlier aptamer-biosensor work showing that cold storage at 4 °C can preserve aptamer-coated sensor function over several weeks^40, 41^. Storage at 4 °C is therefore recommended as the most practical condition, providing reliable performance for at least one month without the added complexity of frozen storage.

### 2.2. 17β-Estradiol electrochemical aptamer-based biosensor development

#### 2.2.1. In vitro characterization for extraction of dose-response and calibration curve

In the next step, the E2 aptamer was modified with a methylene blue (MB) redox reporter to enable reagent-free, single-step electrochemical readout without the need for an exogenous ferri/ferrocyanide redox probe. Square-wave voltammetry (SWV) was selected as the transduction method because it produced a stronger readout than cyclic voltammetry (CV) and differential pulse voltammetry (DPV) under the tested conditions (Figure S4). This choice is also consistent with recent direct comparisons of voltammetric interrogation methods for electrochemical aptamer-based sensors, where SWV provided high signal-to-noise performance and supported reliable sensor interrogation compared with other pulsed voltammetric approaches^42^. To further enhance analytical performance, the effect of electrode nanostructuring on sensitivity was investigated by comparing gold nanoparticle (AuNP)-decorated Au-electrodes with gold nanorod (AuNR)-decorated Au-electrodes. All experiments have been done in binding buffer (2 mM Tris–HCl, pH 7.5, 10 mM NaCl, 0.5 mM KCl, 0.2 mM MgCl₂, 0.1 mM CaCl₂, and 5% ethanol). **Figures 4a,b** show calibration curves obtained with AuNP- and AuNR-modified sensors, respectively. While both nanostructures improve sensitivity relative to bare gold electrodes, the use of AuNRs results in an approximately 20-fold increase in signal response compared to AuNPs^43^. This enhancement is consistent with the higher effective electroactive surface area and geometry-dependent electrochemical activity of nanostructured gold surfaces, which can increase redox-reporter loading, improve electron-transfer efficiency, and enhance the signal-to-noise ratio of electrochemical aptamer-based sensors^44, 45^. The anisotropic morphology of AuNRs may also improve probe accessibility and reduce steric crowding relative to more densely packed nanoparticle films^46^.

**Figure 4.**
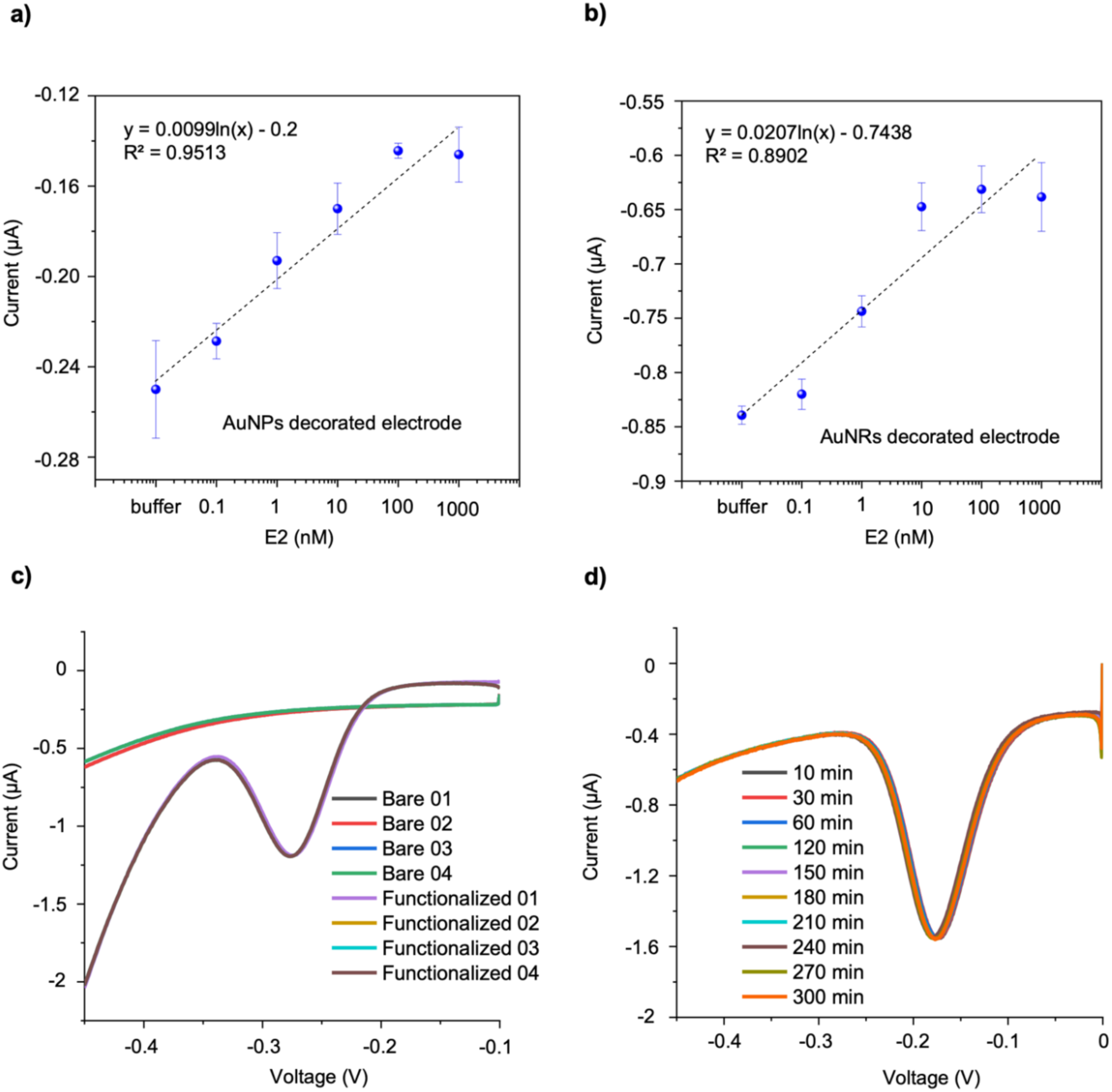
Dose–response and calibration characterization of the electrochemical aptamer-based biosensor (EAB) for estradiol detection using different electrode surface nanostructures. (a) Calibration curve for electrodes decorated with 40 nm citrate-capped AuNPs (*n*=3). (b) Calibration curve for electrodes decorated with 10 nm (diameter) citrate-capped AuNRs (*n*=3). (c) Repeatability of the SWV signal after surface functionalization process for AuNR-decorated electrodes. (d) Stability of the methylene blue peak current for AuNR-decorated sensors over 300 min in binding buffer at room temperature.

Figure 4c demonstrates the reproducibility of the selected functionalization conditions across independently prepared electrodes, indicating consistent MB-tagged aptamer immobilization and reproducible SWV response (*n* = 4, RSD < 1%). Temporal stability of the MB-based assay was evaluated to assess operational stability under the selected buffer and interrogation conditions. As shown in Figure 4d, the electrochemical signal remained stable for up to 300 min, indicating low short-term signal drift and preservation of MB redox activity during the measurement period.

#### 2.2.2. Sensor ex vivo characterization

##### Characterization in artificial sweat and real human sweat (non-pregnant) samples

To assess biosensor performance under more physiologically relevant sample conditions, ex vivo measurements were performed in estradiol-spiked artificial sweat and commercially sourced real human sweat collected from non-pregnant donors (**Figure 5**). In artificial sweat, the device showed a clear concentration-dependent response across the 0.1–1000 nM range, with the measured current becoming progressively less negative as the E2 level increased. The resulting calibration followed a logarithmic relationship, y=0.0148ln (x) – 0.5888, with an *R*^2^ value of 0.9728, indicating reliable behavior in the synthetic matrix. A comparable trend was also observed in real human sweat, where the response was fitted by y=0.0015ln (x) – 0.6709 with the same *R*^2^ value of 0.9728. The lower slope in real sweat compared with artificial sweat indicates approximately one order of magnitude lower signal sensitivity in the native biological matrix. This signal attenuation is likely due to matrix effects arising from endogenous sweat components, ionic-strength differences, nonspecific adsorption, and possible changes in MB redox behavior or aptamer interfacial dynamics^47^. Nevertheless, the preservation of a monotonic logarithmic response over four orders of magnitude in E2 concentration shows that the aptamer-modified sensing interface remains functional in real sweat and can distinguish between different spiked estradiol concentrations despite reduced signal amplitude. In other words, although the biological matrix reduced the overall signal amplitude, it did not compromise the sensor’s ability to distinguish between different estradiol concentrations.

**Figure 5.**
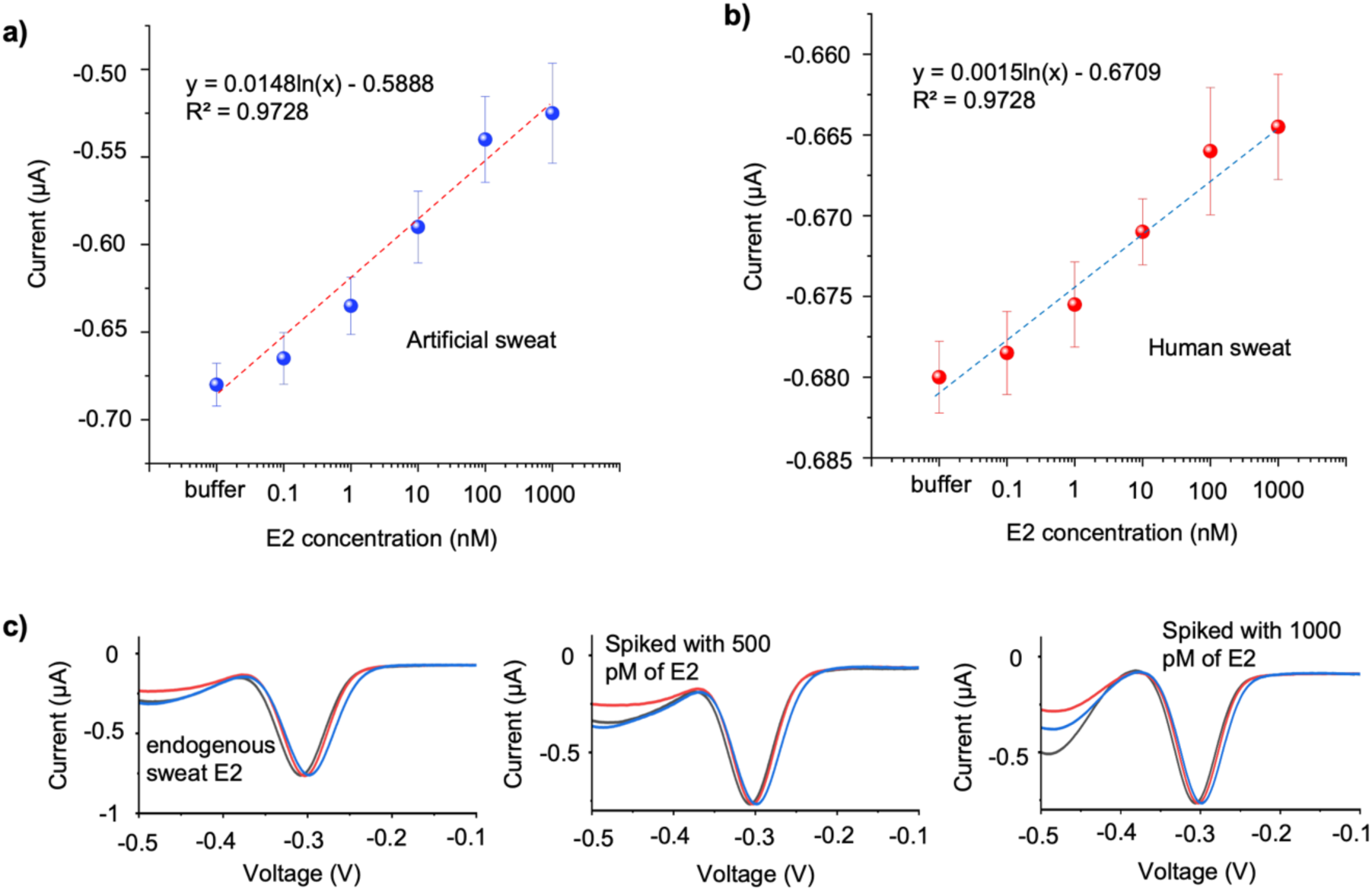
Ex vivo calibration curves of the sensor in response to different concentrations of estradiol spiked into (a) artificial sweat, (b) real human sweat (non-pregnant), and (c) results of spike-and-recovery test with pregnant woman samples before and after spiking.

##### Spike-and-recovery test with pregnant woman sweat samples

Spike-and-recovery analysis was performed by spiking 500 pM and 1000 pM E2 to pregnant-woman sweat samples and quantifying the post-spike E2 concentration using the calibration curve extracted from real human sweat samples (Figure 5b). First, unspiked sweat samples from a woman in the third trimester of pregnancy were directly tested using functionalized sensors to determine the endogenous E2 background level. The SWV response shown in Figure 5c was converted to concentration using the real-sweat calibration equation (Figure 5b), yielding an endogenous sweat E2 concentration of 271.7 ± 23.5 pM. This concentration is physiologically plausible for a third-trimester pregnancy sample because maternal E2 increases progressively across gestation and reaches its highest levels during late pregnancy, with longitudinal pregnancy studies reporting markedly elevated circulating E2 concentrations in the third trimester^48, 49^. In addition, recent wearable sweat-sensing studies have demonstrated pM-level E2 detection in sweat and reported correlation between sweat and blood E2 profiles, supporting the feasibility of detecting estradiol in this concentration range in sweat^19^. However, because pregnancy-specific sweat E2 reference intervals have not yet been established and estrogen concentrations show large interindividual variability during pregnancy, this value should be interpreted as a matrix-calibrated endogenous sweat E2 concentration from a third-trimester sample rather than as a clinical reference value.

The same procedure was then performed for sweat samples spiked with 500 and 1000 pM E2. The measured SWV responses corresponded to the calculated E2 concentrations of 854.5 ± 54.1 pM and 1403.3 ± 64.6 pM, respectively. The expected post-spike concentrations, calculated as endogenous E2 plus the added spike, were approximately 771.7 pM for the 500 pM spike and 1271.7 pM for the 1000 pM spike. The recovery values were 116.6 ± 6.1% and 113.2 ± 4.1% for the 500 pM and 1000 pM spike levels, respectively. The absolute percentage errors relative to the expected post-spike concentrations were 10.66 ± 3.65% and 10.32 ± 3.04%, respectively. These values fall within the commonly used acceptable recovery range for bioanalytical and biosensor matrix studies, supporting reliable quantification of E2 in pregnant-woman sweat despite modest positive bias^50, 51^. The recovery values above 100% suggest a mild matrix-enhancement effect or calibration-transfer bias when applying the non-pregnant real-sweat calibration curve to pregnant-woman sweat. Nevertheless, the consistency between the two spike levels indicates that the sensor preserves quantitative response behavior in the pregnant-sweat matrix. These results support the feasibility of the MB-labeled E2 aptasensor for ex vivo estradiol quantification in pregnant-woman sweat.

Having established the biosensor’s in vitro and ex vivo analytical performance, we next examine flow-programmed self-regeneration to determine whether the same sensing interface could be restored for repeated E2 measurements, a key requirement for reusable and longitudinal sweat monitoring.

### 2.3. Flow-programmed self-regeneration and dynamic operation of the estradiol EAB sensor

To examine dynamic-flow experiments and flow-programmed self-regeneration of estradiol EAB sensor, an experimental setup was developed as described in SI. Sensor functionalization and preparation was performed based on developed protocols in previous sections and as described in SI. Experimental results are summarized in **Figure 6**. In these dynamic-flow experiments, the developed biosensor was integrated into the flow chamber and interrogated by SWV while the inlet solution was sequentially switched between blank binding buffer and E2-containing buffer. First, a single buffer/1 nM E2/buffer cycle was performed using 60 min steps to evaluate whether the sensor response could be reversed after target exposure. At a constant 2 µL min⁻¹ flow rate, E2 introduction produced a clear current shift from the buffer baseline, and subsequent blank-buffer perfusion drove the signal back toward its initial level, demonstrating reversible E2 recognition and partial regeneration of the sensing interface (Figure 6a). When the E2 injection was kept at 2 µL min⁻¹ but the buffer-wash flow was increased to 8 µL min⁻¹, the recovery phase became sharper, and the baseline was restored more efficiently (Figure 6b). The same trend was observed during sequential exposure to increasing E2 concentrations from 10 pM to 100 nM, as the sensor produced concentration-ordered current responses, while higher buffer-wash flow reduced baseline carryover between pulses and improved repeatability of the regenerated state (Figure 6c,d). These results show that the sensing interface remains electrochemically active during repeated target/wash cycles and that buffer-flow rate and washing duration are key operational parameters controlling apparent regeneration efficiency.

**Figure 6.**
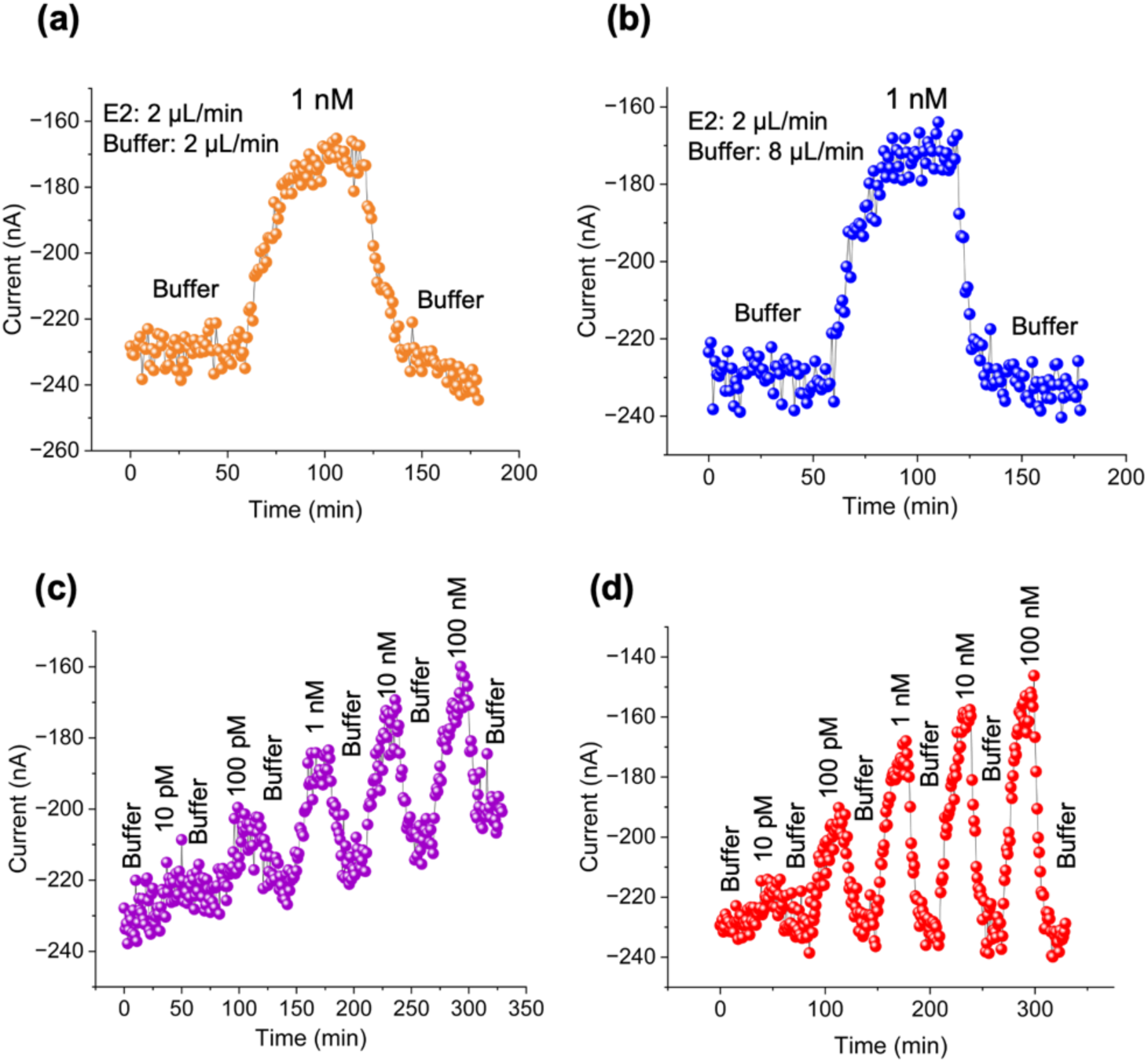
Dynamic flow characterization of the estradiol EAB sensor at different flow-rate conditions. (a) Dynamic response during sequential injection of buffer–1 nM estradiol–buffer, 60 min per step, at 2 µL min⁻¹. (b) Dynamic response during one buffer–1 nM estradiol–buffer cycle, 60 min per step, at 8 µL min⁻¹ (buffer) and 2 µL min⁻¹ (estradiol). (c) Dynamic response during sequential injection of buffer and different estradiol concentrations, 30 min per step, at 2 µL min⁻¹ for both buffer and estradiol solutions. (d) Dynamic response during sequential injection of buffer and different estradiol concentrations, 30 min per step, at 8 µL min⁻¹ for buffer and 2 µL min⁻¹ for estradiol solutions.

Based on these results, the estradiol EAB sensor can be regenerated by flow programming alone, by switching the inlet from estradiol-containing buffer to blank buffer, without any chemical stripping step. The reversible interaction between the surface-confined aptamer and estradiol can be described as Equation (1):

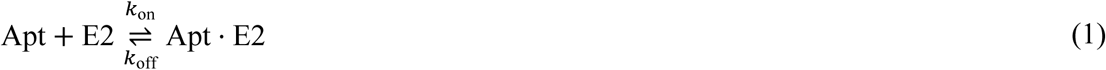

The kinetic and thermodynamic parameters are related by Equation (2):

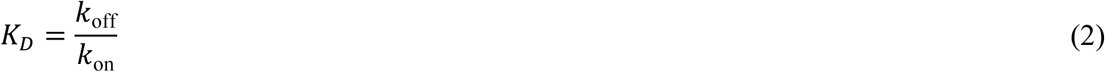

where *k*_on_ is the association rate constant, *k*_off_ is the dissociation rate constant, and *K_D_* is the dissociation constant. For a surface-confined aptamer layer, the fractional occupancy of bound aptamers, *θ*, follows the standard surface-binding kinetic expression shown in Equation (3):

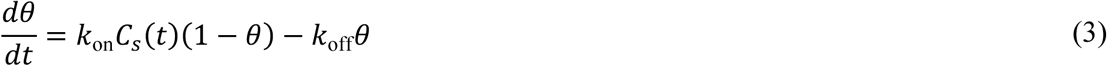

In this equation, *C_s_* (*t*) is the local free estradiol concentration immediately above the aptamer layer, rather than simply the inlet concentration. This distinction is critical for interpreting Figure 6 because, in flow-based surface biosensors, the sensor response is governed by the coupled effects of convection, diffusion, and surface reaction. Thus, regeneration depends not only on molecular dissociation from the aptamer but also on how rapidly free estradiol is removed from the chamber and the near-electrode diffusion layer. This treatment follows the convection-reaction-diffusion framework used for surface-based biosensors and microfluidic surface-reaction systems^52^.

Under a constant interfacial estradiol concentration, Equation (3) gives the Langmuir equilibrium occupancy in Equation (4):

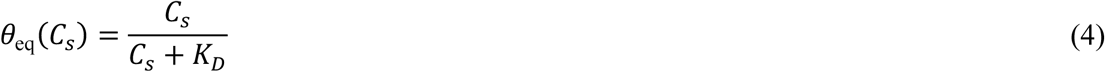

For a heterogeneous immobilized aptamer population, the response can be described by an apparent Hill-Langmuir form, Equation (5):

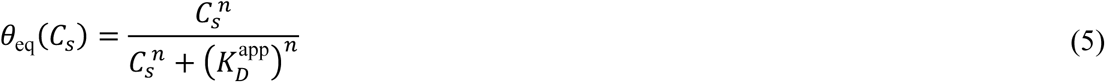

where *K_D_^app^* is the apparent dissociation constant of the surface-confined aptamer layer and *n* accounts phenomenologically for surface heterogeneity, steric effects, non-uniform probe orientation, or other non-idealities. This framework explains the monotonic increase in current response from the picomolar to nanomolar regime in Figure 6c,d and the tendency toward a plateau at higher concentrations, as a finite population of surface-confined aptamers becomes progressively occupied as estradiol concentration increases.

For a target pulse of duration *t_p_*, Equation (3) gives the transient association response in Equation (6):

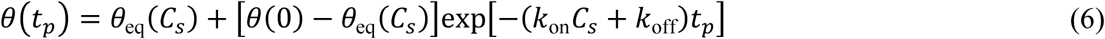

This expression indicates that the response during estradiol injection approaches a quasi-steady value with an apparent time constant controlled by both association, *k*_on_*C_s_*, and dissociation, *k*_off_. During an ideal buffer wash in which *C_s_* → 0, the rebinding term becomes negligible and Equation (3) reduces to Equation (7):

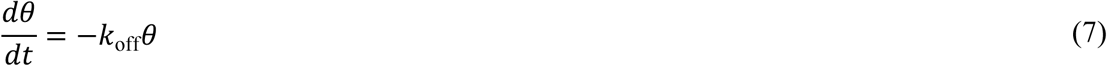

The corresponding solution is Equation (8):

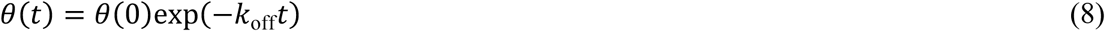

Therefore, the theoretical requirement for passive regeneration is not that flow mechanically breaks the estradiol-aptamer complex, but that flow drives the local free estradiol concentration, *C_s_*, toward zero. Once *C_s_* is minimized, rebinding is suppressed and the surface occupancy decays through the normal thermal dissociation pathway of the E2-aptamer complex. The corresponding dissociation half-life is given by Equation (9):

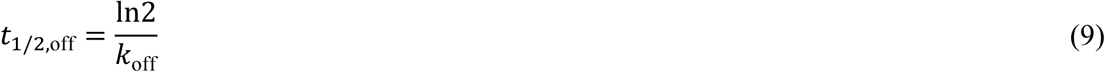

Thus, the measured recovery time in Figure 6 should be interpreted as an apparent regeneration time that combines chamber exchange, near-surface mass transfer, rebinding suppression, intrinsic aptamer dissociation, and electrochemical signal relaxation.

The single-cycle experiments in Figure 6a,b are consistent with this model. In both cases, introduction of 1 nM estradiol produced a pronounced and reversible current shift from the initial buffer baseline to a quasi-steady plateau within tens of minutes, confirming target recognition under continuous flow. Since the estradiol step was maintained at 2 µL min⁻¹ in both panels, the comparable 1 nM response amplitudes indicate that analyte delivery during the sensing step was sufficient to establish similar target occupancy. When the influent was returned to blank buffer, the current relaxed back toward its initial baseline, indicating dissociation of the estradiol-aptamer complex and restoration of the unbound interfacial state of the probe layer. The sharper recovery in Figure 6b, where the buffer flow was increased to 8 µL min⁻¹, indicates that apparent regeneration is accelerated when residual estradiol is removed more rapidly from the chamber and near-electrode region.

This behavior can be quantified using a simple chamber mass balance. If *C_b_* (*t*) is the bulk estradiol concentration in the flow chamber, *Q* is the volumetric flow rate, *V*_ch_ is the chamber volume, *A*_WE_ is the active working-electrode area, and *Γ_T_* is the surface density of active aptamer probes, then the chamber balance is formulated as Equation (10) ^53–56^:

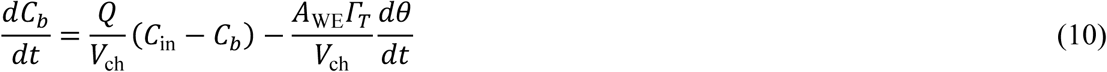

During buffer washing, *C*_in_ = 0. If surface release and uptake are small relative to the bulk chamber inventory, Equation (10) reduces to the exponential washout relation in Equation (11):

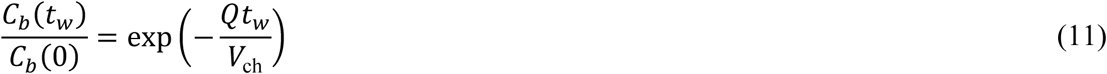

Where *t*_w_ is the wash duration. Using the actual circular chamber geometry of this study (diameter of 9 mm and height of 0.6 mm), the chamber volume is calculated as *V_ch_* = 38.17 µL, and the working-electrode is circular with diameter of 4 mm and area of *A_WE_* = 0.1257 cm^2^. For the 60 min buffer wash in the single-cycle experiments (Figure 6a,b), Equation (11) predicts that a 2 µL min⁻¹ wash exchanges *Qt_w_/V_ch_* = 3.14 chamber volumes and leaves the residual fraction calculated as 4.31 × 10^-2^. This corresponds to approximately 4.3% of the previous bulk estradiol concentration under an ideal well-mixed approximation. In contrast, a 60 min wash at 8 µL min⁻¹ exchanges 12.57 chamber volumes and leaves the residual fraction of 3.46 × 10^-6^. This corresponds to approximately 0.00035% of the previous bulk estradiol concentration. Therefore, the higher buffer flow is expected to remove free estradiol much more efficiently, explaining the sharper decay and improved baseline restoration observed in Figure 7b.

**Figure 7.**
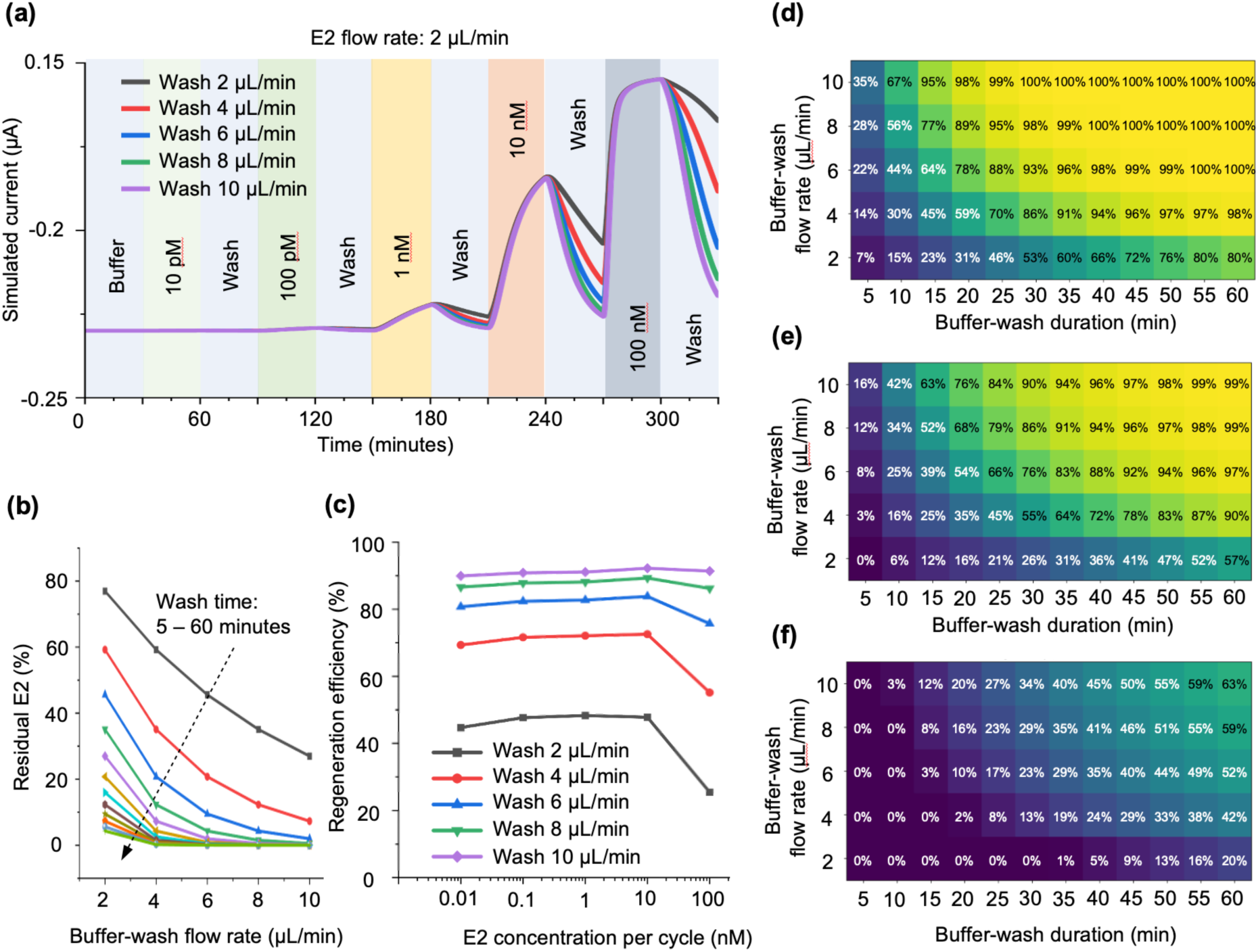
Simulation-guided design space for flow-programmed regeneration of the estradiol EAB sensor. (a) Simulated cyclic SWV current response to sequential E2 pulses of 10 pM, 100 pM, 1 nM, 10 nM, and 100 nM, with 30 min E2 exposure at 2 µL min⁻¹ and intermediate buffer-wash steps at 2, 4, 6, 8, and 10 µL min⁻¹. (b) Predicted residual bulk E2 remaining after buffer washing as a function of wash-flow rate and wash duration. (c) Regeneration efficiency extracted from the coupled binding–transport simulation for each E2 concentration and wash-flow condition. (d–f) Regeneration-efficiency maps showing the minimum recovery across the full 10 pM–100 nM E2 sequence as a function of buffer-wash flow rate and wash duration for representative apparent dissociation constants of (d) 20 nM, (e) 5 nM, and (f) 1 nM. All simulations used the experimental chamber volume of 38.17 µL, working-electrode area of 12.57 mm², and E2 injection flow of 2 µL min⁻¹. (Python code and related parameters are in Supporting Information).

The concentration-sequence experiments in Figure 6c,d further clarify the role of flow rate in self-regeneration. Under the symmetric low-flow condition of 2 µL min⁻¹ for both estradiol and buffer (Figure 6c), sequential injections of 10 pM, 100 pM, 1 nM, 10 nM, and 100 nM generated stepwise current changes with the expected concentration ordering. However, the intermediate buffer segments did not fully restore the original baseline, and a progressive baseline shift was observed as the sequence advanced, particularly after the higher-concentration pulses. This behavior indicates incomplete removal of estradiol from the chamber and near-electrode region, allowing residual interfacial estradiol to maintain a nonzero rebinding term, *K_on_ C_s_* (1-*θ*), during the wash step.

For the 30 min wash intervals used in Figure 6c,d, the impact of flow rate is even more evident. At 2 µL min⁻¹, the residual fraction is calculated as 0.208. This means that approximately 20.8% of the previous bulk estradiol concentration may remain after each 30 min wash under the ideal well-mixed approximation. At 8 µL min⁻¹, the residual fraction is calculated as 1.86 × 10^-3^. This means that only approximately 0.186% remains. Thus, increasing the buffer flow from 2 to 8 µL min⁻¹ decreases the residual bulk estradiol concentration by approximately 112-fold for a 30 min wash. This theoretical washout ratio directly supports the experimental observation that Figure 6d exhibits much lower baseline carryover than Figure 6c. The 8 µL min⁻¹ wash decreases *C*_s_ (*t*), suppresses rebinding, and allows dissociation to dominate the regeneration phase.

### 2.4. Theoretical validation of flow-programmed regeneration and design-space mapping

To support the experimental flow-regeneration results and define the operating space of the device, a numerical model was developed from the coupled surface-binding and chamber-transport equations described above (Python code and related parameters are in Supporting Information). The simulation used the actual device geometry, including a circular flow chamber with *d_ch_* = 9mm, *h* = 0.6 mm, and *V_ch_* = 38.17 µL, together with a circular working electrode with *d_WE_* = 4 mm, and *A_WE_* = 12.57 mm^2^. The estradiol injection flow was fixed at 2 µL min⁻¹, while the buffer-wash flow was varied from 2 to 10 µL min⁻¹. As shown in **Figure 7a**, the simulated cyclic traces reproduce the major experimental features observed in Figure 6c,d, including stepwise concentration-dependent current modulation, incomplete baseline recovery at low wash flow, and improved regeneration as the wash-flow rate increases. This agreement supports the interpretation that regeneration is governed by the coupled effects of chamber exchange, interfacial analyte depletion, suppression of rebinding, and reversible recovery of the aptamer layer during the buffer phase^52, 57^.

The analyses in Figure 7b,c establish the physical link between chamber-scale analyte clearance and functional recovery of the EAB interface. Figure 7b first isolates the geometric washout process, showing that the residual free estradiol concentration decreases according to *C_b_*(*t*_w_)/*C_b_*(0) = exp(−*Qt*_w_/*V*_ch_). This relationship is important because it converts the flow chamber geometry into a quantitative regeneration constraint, as for the present chamber volume of 38.17 µL, a 30 min wash at 2 µL min⁻¹ exchanges only 1.57 chamber volumes and can leave approximately 20.8% of the previous free estradiol in the chamber, whereas the same wash duration at 8 µL min⁻¹ exchanges 6.29 chamber volumes and lowers the residual fraction to approximately 0.186%. Thus, increasing the wash flow from 2 to 8 µL min⁻¹ produces an approximately 112-fold reduction in residual free estradiol during a 30 min wash. Figure 7c then extends this geometric washout calculation into the full coupled binding–transport model, where residual *C_b_* (*t*) maintains a nonzero interfacial concentration *Cs* (*t*), promotes rebinding, and limits return of the aptamer layer to its pre-injection state. The resulting regeneration-efficiency trends show that low-flow washing is not simply slower, it leaves the sensor in a history-dependent state with measurable residual signal after each pulse. In contrast, higher wash-flow rates reduce the local analyte burden, suppress carryover, and restore the sensor closer to a common baseline between measurements. Together, Figure 7b,c transform the experimental observation of improved recovery at higher buffer flow into a quantitative design rule for selecting wash-flow rate and wash duration in continuous EAB operation.

Figures 7d–f extend this framework by mapping regeneration efficiency as a function of buffer-wash flow rate and wash duration for three representative aptamer affinity regimes. These heatmaps were generated using the same concentration sequence used experimentally and in Figure 7a (including 10 pM, 100 pM, 1 nM, 10 nM, and 100 nM E2), with each E2 pulse fixed at 30 min and 2 µL min⁻¹. The color scale reports the minimum regeneration efficiency across the full sequence, providing a conservative measure of the worst-case recovery expected under each washing condition. The maps show that regeneration becomes progressively more demanding as the apparent affinity increases. The *K_D_* = 20 nM case can be efficiently regenerated over a broad flow–duration window, the *K_D_* = 5 nM case requires higher flow and/or longer washing, and the *K_D_* = 1 nM case remains difficult to fully regenerate even under stronger wash conditions. This affinity dependence is important because it identifies the boundary of buffer-only regeneration, as for very high-affinity receptors, particularly sub-nM systems, simple washing may fall outside the practical regeneration window even at 10 µL min⁻¹ for 60 min, and additional reset strategies may be required. Together, Figure 7 establishes flow-programmed regeneration as a quantitatively designable process governed by chamber volume, wash-flow rate, wash duration, and aptamer binding kinetics, providing a practical framework for integrating reagent-free EAB regeneration into wearable-compatible and automated biosensing systems.

## 3. Conclusion

This work demonstrates a flow-triggered, self-regenerative electrochemical aptamer-based platform for repeated 17β-estradiol sensing with a potential application in continuous E2 sensing from sweat. By coupling target recognition and interface recovery to buffer-flow programming, the device addresses a key limitation of continuous affinity biosensing. The optimized platform combines picomolar estradiol detection, selective response against tested interferents, AuNR-enhanced electrochemical readout, and stable methylene-blue signal behavior during extended operation. Importantly, the sensor retained concentration-dependent performance in artificial sweat and real human sweat. Furthermore, spike-and-recovery experiments in pregnant woman sweat samples further support its potential for noninvasive hormone monitoring in complex biological matrices.

Dynamic-flow experiments and coupled binding–transport simulations show that regeneration is governed not only by aptamer–target dissociation, but also by chamber exchange, near-surface analyte depletion, and suppression of rebinding during the buffer-wash phase. Increasing the buffer-wash flow rate reduced residual estradiol, minimized carryover, and improved baseline recovery, while the simulation framework translated these trends into practical design rules linking chamber volume, wash-flow rate, wash duration, and apparent aptamer affinity. Although further validation is required for long-term on-body operation, sweat-rate variability, interindividual calibration, and fully integrated wearable deployment, the present study establishes buffer-flow programming as a simple, reagent-free, and wearable-compatible strategy for regenerable electrochemical aptamer sensing, with broader applicability to continuous affinity biosensors limited by saturation, hysteresis, and slow recovery.

## Acknowledgements

The authors acknowledge Ata Golparvar from Delft University of Technology and Gian Luca Barbruni from École Polytechnique Fédérale de Lausanne for useful discussions. The research has been financially supported by the Innosuisse grant “*MatPacth*” (project # 116.366 IP-ENG) and by the SNF Bridge proof-of-concept grant “*HormoTrack*” (project # 40B1-0_229802 / 2). The research has been also supported by internal EPFL funding.

## Conflict of Interest

All authors declare no financial/commercial conflicts of interest.

Received: ((will be filled in by the editorial staff)) Revised: ((will be filled in by the editorial staff)) Published online: ((will be filled in by the editorial staff))

## Supporting Information

Supporting Information is available from the Wiley Online Library.

