## Supplementary material for "Flow-Triggered Regeneration of Electrochemical Aptasensor for Continuous Monitoring of 17β-Estradiol": SI file

#### Methods

##### 1. Chemicals and Materials

The anti-17β-estradiol thiol-modified DNA aptamer was purchased from Creative Biolabs (Cat. No. CTApt-2054). 17β-Estradiol was obtained from EMD Millipore/Merck (Cat. No. 3301-1GM). Estriol (Cat. No. E1253-500MG), progesterone (Cat. No. P8783-1G), and testosterone (Cat. No. 46923-250MG-R) were used for selectivity studies and were purchased from Sigma-Aldrich. Citrate-stabilized gold nanoparticles (AuNPs) with diameters of 40 nm (Cat. No. 742007-25ML, OD 1) were purchased from Sigma-Aldrich/MilliporeSigma. Gold nanorods (Cat. No. 900364-25ML, 10 nm diameter, $\lambda_{max}$ = 980 nm, dispersed in $H_2O$ containing CTAB) were also obtained from Sigma-Aldrich. Cysteamine hydrochloride (Cat. No. M6500-25G, ≥98%), 6-mercapto-1-hexanol (MCH; Cat. No. 451088-5ML, 97%), and tris(2-carboxyethyl)phosphine hydrochloride (TCEP; Cat. No. C4706-10G) were purchased from Sigma-Aldrich and used for surface modification, blocking, and reduction of the disulfide-protected thiolated aptamer, respectively. Trizma hydrochloride solution (1 M, pH 7.5, BioPerformance certified; Cat. No. T2319-100ML), sodium chloride (Cat. No. S9888-25G, ACS reagent, ≥99.0%), potassium chloride (Cat. No. P5405-250G), calcium chloride (Cat. No. C4901-100G, anhydrous, ≥97%), and magnesium chloride (Cat. No. M8266-100G, anhydrous, ≥98%) were used for preparation of the binding and measurement buffers. Potassium ferricyanide(III) (potassium hexacyanoferrate(III); Cat. No. 702587-250G, 99%) was used for electrochemical characterization. Estradiol ELISA Kit (Item No. Cay501890) was purchased from Cayman Chemical. Ethanol (Cat. No. 493546-1L, 200 proof) and nuclease-free water (Cat. No. W4502-50ML) were purchased from Sigma-Aldrich. All chemicals were used as received without further purification unless otherwise stated. Electrochemical measurements were performed using a MultiEmStat4 LR potentiostat/galvanostat (PalmSens, The Netherlands). Dynamic microfluidic flow experiments were conducted using Fluigent FLOW-EZ pressure controller modules integrated with a LINK-EZ interface (Fluigent, France). Aptamer preparation and incubation steps were supported by a ThermoMixer C equipped with a ThermoTop heated lid (Eppendorf, Germany), a Z 300 K centrifuge (Hermle, Germany), a VWR analog vortex mixer, and an ML104T analytical balance (Mettler Toledo, Switzerland). Microfluidic housing for dynamic experiments was produced using a Form 4 3D printer (Formlabs). Real human sweat samples and pregnant woman sweats sample (donor number: T6820, age: 27, trimester: third trimester) were purchased from Medix biochemical (USA, Lot: 24-01-613-T6820).

#### 2. Electrochemical cleaning and activation of the gold biosensor surface

Prior to aptamer immobilization, the gold working electrode surface was electrochemically cleaned and activated in 0.5 M $H_2SO_4$ to remove residual organic and inorganic contaminants and to generate a reproducible gold surface. The electrode was immersed in 5 mL of freshly prepared 0.5 M $H_2SO_4$ together with an Ag/AgCl reference electrode and a suitable counter electrode. All potentials were applied versus Ag/AgCl.

The cleaning procedure was initiated by applying a short anodic/cathodic potential sequence. The gold electrode was first held at +2.0 V for 5 s, followed immediately by a cathodic hold at −0.35 V for 10 s. This potential-pulse step was used to oxidize and reduce surface species and remove loosely adsorbed impurities from the gold surface.

After this initial electrochemical polishing step, the electrode was further cleaned by repeated cyclic voltammetry in 0.5 M $H_2SO_4$. Twenty consecutive CV scans were recorded in 5 mL of 0.5 M $H_2SO_4$ over the potential window of −0.30 to +1.55 V vs Ag/AgCl at a scan rate of 4 V $s^{-1}$, using a sample interval of 0.001 V. These rapid scans were performed to promote repeated gold surface oxidation and reduction, thereby removing residual contaminants and activating the electrode surface.

Following the high-scan-rate cleaning cycles, the electrode was transferred to a fresh 0.5 M $H_2SO_4$ solution and subjected to an additional conditioning step. Four CV scans were recorded over the same potential window, −0.30 to +1.55 V vs Ag/AgCl, at a slower scan rate of 0.1 V $s^{-1}$ with a sample interval of 0.001 V. The slower final scans were used to confirm the characteristic voltammetric response of a clean gold surface, including gold oxide formation during the anodic sweep and oxide reduction during the reverse scan.

After completion of the cleaning process, the electrode was thoroughly rinsed with deionized water to remove residual sulfuric acid and dried gently under nitrogen or clean air. The cleaned gold surface was then used immediately for subsequent biosensor functionalization to minimize adventitious contamination.

#### 3. Preparation of 17β-Estradiol biosensor

Planar gold electrodes were used as the transducer platform for fabrication of the 17β-estradiol biosensor. The electrodes were electrochemically cleaned in 0.5 M $H_2SO_4$ by cyclic voltammetry from -0.2 to +1.5 V versus Ag/AgCl at 100 mV $s^{-1}$ for 10 cycles, followed by thorough rinsing with deionized water. To generate a nanostructured interface with enhanced electroactive area, the cleaned electrodes were immersed overnight in 10 mM cysteamine hydrochloride to form an amine-terminated layer and subsequently incubated overnight in citrate-capped AuNP/AuNR solution (40 nm average diameter). The resulting AuNP/AuNR-coated electrodes were dried and heated at 60 °C for 1 h to improve nanoparticle adhesion and surface stability. A thiol-modified anti-17β-estradiol DNA aptamer was reconstituted in deionized water, aliquoted, and stored at -20 °C until use. Before immobilization, the aptamer was diluted in binding buffer, thermally denatured at 95 °C for 5 min, and allowed to refold at room temperature for 15 min. To generate free terminal thiol groups for surface anchoring, the aptamer was treated with a 100-fold molar excess of TCEP for 2 h at room temperature. Subsequently, 40 μL of 100 nM aptamer solution was deposited onto each AuNP/AuNR-modified electrode and incubated overnight in a humidified chamber at room temperature, enabling self-assembly of the aptamer on the gold surface via Au–S interactions. After immobilization, the electrodes were backfilled with 1 mM MCH in PBS for 2 h to block unreacted gold sites and reduce nonspecific binding. The completed biosensor was rinsed thoroughly with deionized water and used immediately for analytical measurements.

#### 4. In vitro EIS characterization of biosensor

In vitro electrochemical impedance spectroscopy (EIS) characterization of the 17β-estradiol biosensor was performed using a MultiEmStat4 LR potentiostat/galvanostat (PalmSens) in a three-electrode configuration, with the functionalized planar gold biosensor serving as the working electrode, an Ag/AgCl electrode as the reference electrode, and an pt electrode as the counter electrode. The biosensor surface was functionalized according to the procedure described in Section 4.2. For the binding assay, the aptamer-functionalized electrodes were incubated with 17β-estradiol solutions prepared in binding buffer consisting of 2 mM Tris–HCl (pH 7.5), 10 mM NaCl, 0.5 mM KCl, 0.2 mM MgCl2, 0.1 mM CaCl2, and 5% ethanol. After the binding step, EIS measurements were carried out in 5 mM K3 [Fe(CN)6] prepared in PBS (pH 7.4). A frequency sweep from 1 Hz to 100 kHz was applied with an amplitude of 50 mV. The resulting impedance spectra were recorded as Nyquist plots and fitted with an appropriate equivalent-circuit model to extract the charge-transfer resistance ($R_{\mathrm{ct}}$). The extracted $R_{\mathrm{ct}}$value was used as the analytical output parameter of the biosensor.

#### 5. In vitro SWV characterization of biosensor

In vitro square-wave voltammetry (SWV) characterization of the 17β-estradiol biosensor was performed using a MultiEmStat4 LR potentiostat/galvanostat (PalmSens). The aptamer-functionalized planar gold biosensor served as the working electrode, while the integrated on-sensor reference and counter electrodes were used for electrochemical measurements. The biosensor surface was prepared according to the functionalization procedure. For target binding, the functionalized sensors were incubated with 17β-estradiol solutions prepared in binding buffer consisting of 2 mM Tris–HCl (pH 7.5), 10 mM NaCl, 0.5 mM KCl, 0.2 mM $MgCl_2$, 0.1 mM $CaCl_2$, and 5% ethanol. Following incubation, SWV measurements were carried out in binding buffer. The voltammograms were recorded from 0 to -0.45V using an amplitude of 50 mV and a frequency of 50 Hz. The resulting peak current response was used as the analytical output of the biosensor.

#### 6. Preparation of artificial sweat

Artificial sweat was prepared to mimic key ionic and metabolic components of human sweat. The solution contained 5.5 mM lactic acid, 3 mM $NH_4^+$, 22 mM urea, 50 µM $Mg^{2+}$, 0.4 mM $Ca^{2+}$, 25 µM uric acid, and E2 concentrations ranging from 0.1 to 1000 nM. The formulation was prepared at the pH of 7.5 and samples were used immediately after preparation.

#### 7. Dynamic flow-programmed regeneration experiments using Fluigent microfluidics and SWV readout

Dynamic flow-regeneration experiments were performed using a Fluigent pressure-driven microfluidic setup integrated with SWV readout from a MultiEmStat4 multi-channel potentiostat/galvanostat/impedance analyzer. Blank binding buffer and estradiol-containing binding buffer were loaded into separate reservoirs and delivered to the sensor through Fluigent Flow EZ pressure/flow controller modules. Fluid selection between buffer and E2 solutions was performed using Fluigent switching valves, including a 2-SWITCH valve for buffer/E2 exchange and an M-SWITCH valve for sequential delivery of multiple E2 concentrations. Before each experiment, the tubing, valves, and flow chamber were primed with binding buffer to remove air bubbles, stabilize fluidic resistance, and establish a stable electrochemical baseline. The functionalized EAB sensor was mounted in a custom flow chamber with a circular chamber diameter of 9 mm and height of 0.6 mm, corresponding to a geometric chamber volume of 38.17 µL. The circular working electrode had a diameter of 4 mm and an active geometric area of 12.57 $mm^2$. The sensor was connected to the MultiEmStat4 in a three-electrode configuration, and SWV was recorded in binding buffer from 0 to −0.45 V using a 50 mV amplitude and 50 Hz frequency. The methylene-blue peak current was extracted from each SWV scan and plotted over time to follow E2 binding and buffer-induced regeneration.

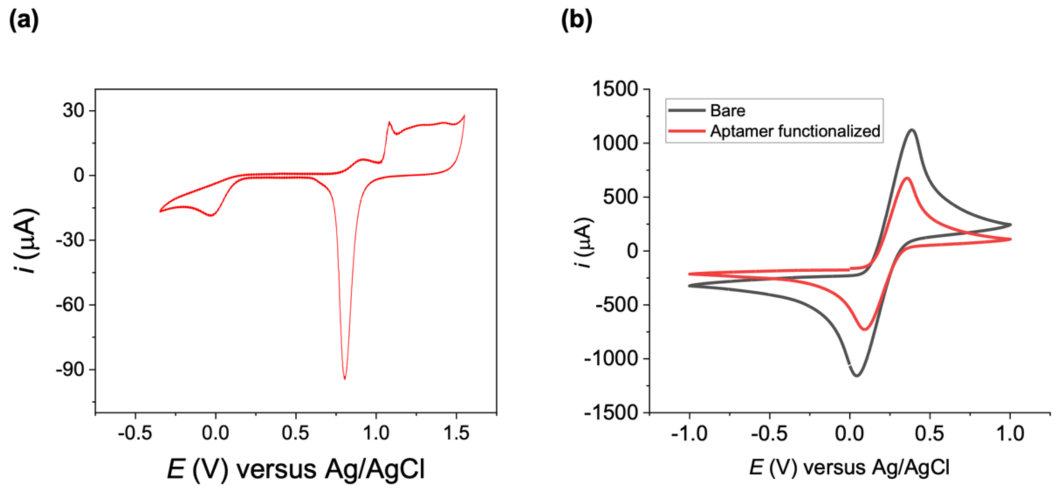


**Figure S1**. Electrochemical characterization of gold electrode cleaning and aptamer functionalization. (a) Measured cyclic voltammogram of a clean gold electrode recorded in 0.5 M $H_2SO_4$, showing the characteristic gold surface oxidation and reduction features after electrochemical cleaning. (b) Cyclic voltammograms of the sensor before and after aptamer functionalization, recorded using the ferri/ferrocyanide redox couple, $[Fe(CN)_6]^{3-/4-}$. Compared with the bare electrode, the aptamer-functionalized electrode shows reduced redox peak currents and altered peak separation, consistent with formation of an aptamer layer on the electrode surface that partially hinders interfacial electron transfer.

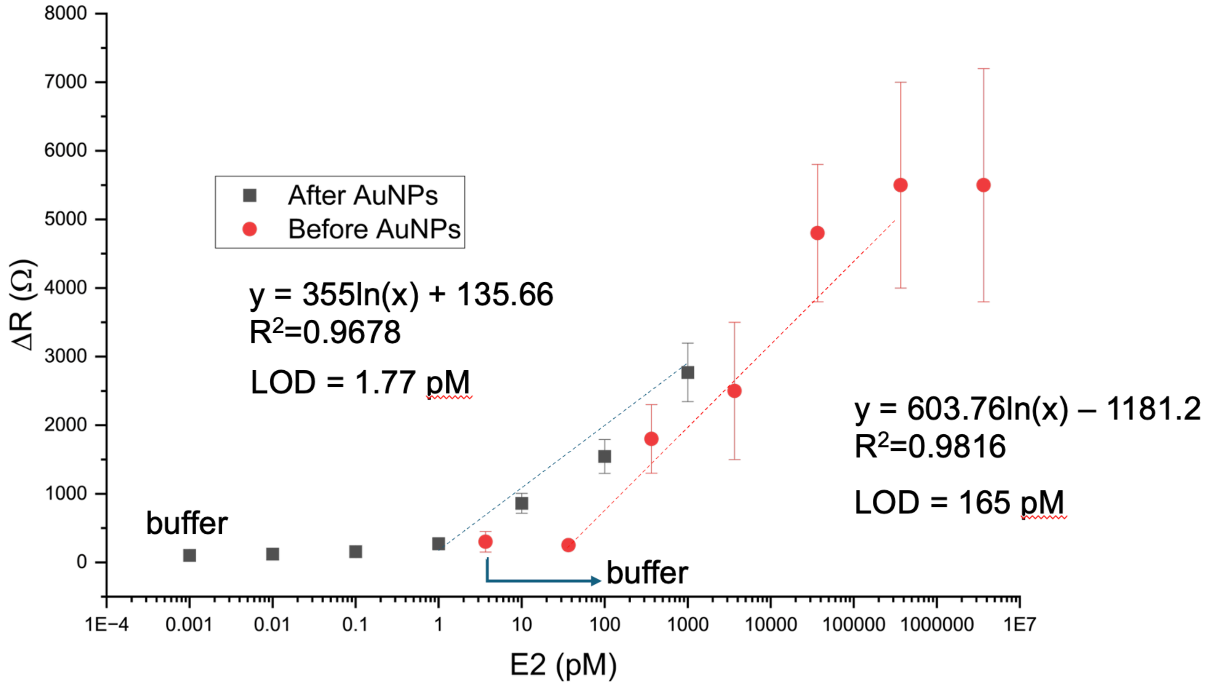


**Figure S2.** Comparative study of the effect of AuNP deposition on the bare electrode in improving the sensor's LOD for E2 detection and quantification.

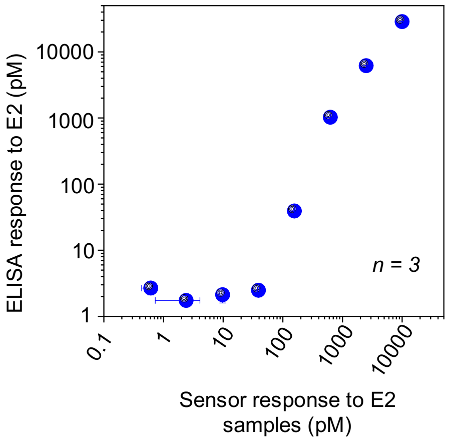


**Figure S3.** Analytical performance benchmarking of the sensor against commercial ELISA assay using matched E2 solutions.

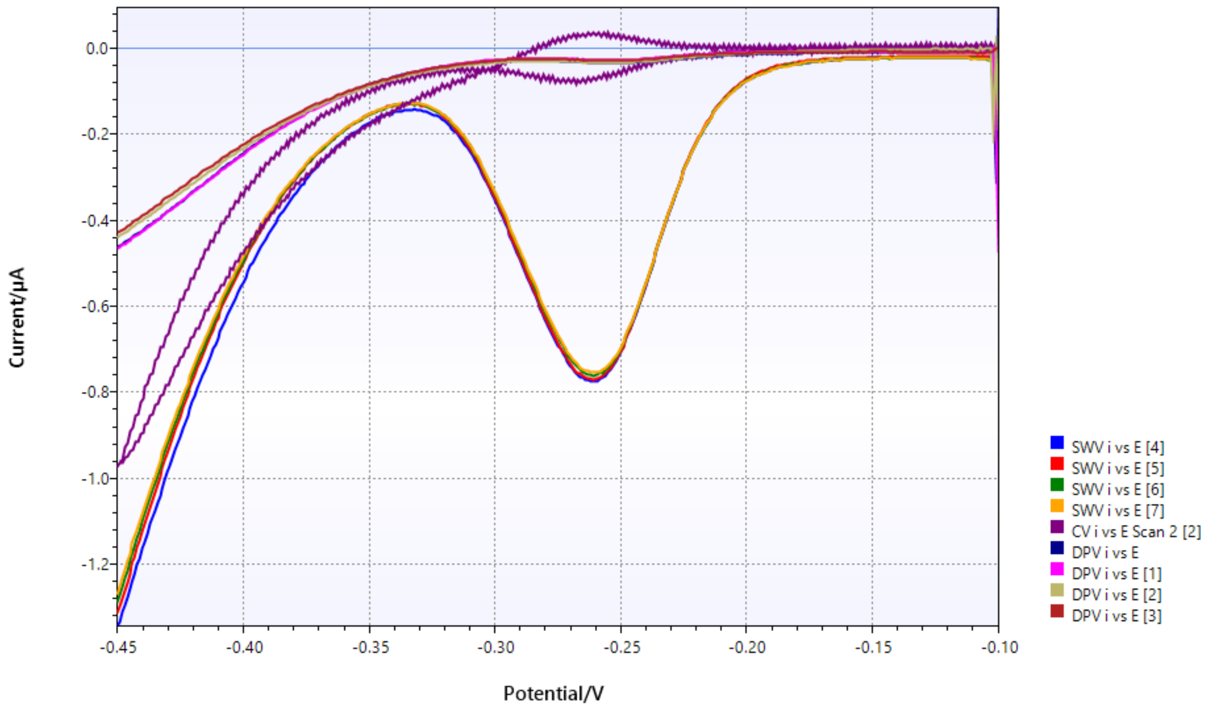


**Figure S4.** Comparison of different measurement techniques of CWV, CV and DPV. As the experiment indicates, the SWV provides strongest response to the functionalized E2 biosensor.

### 8. Developed code for numerical modeling of developed E2 biosensor (Figure 7)

```
"""
Figure 8a simulation code:
Simulated cyclic SWV current response of the developed estradiol EAB sensor during repeated
E2 injection/wash cycles at different buffer-wash flow rates.

Model components:
1) Chamber exchange:
   dCb/dt = (Q/Vch)(Cin - Cb) - (A_WE*Gamma_T/Vch)(dtheta/dt)

2) Interfacial mass transfer:
   km(Cb - Cs) = Gamma_T [kon*Cs(1 - theta) - koff*theta]

3) Surface occupancy:
   dtheta/dt = kon*Cs(1 - theta) - koff*theta

4) Current mapping:
   I(t) = I_unbound + DeltaI*theta(t)

Protocol:
- E2 sequence: 10 pM, 100 pM, 1 nM, 10 nM, 100 nM
- E2 pulse: 30 min at 2 uL/min
- Wash step: 30 min at 2, 4, 6, 8, or 10 uL/min
- Initial buffer: 30 min
"""

from pathlib import Path
import csv
import math
import numpy as np
import matplotlib.pyplot as plt


# ==============================================================================
# Protocol of E2 dose injection and duration
# ==============================================================================

E2_CONCENTRATIONS_M = np.array([
    10e-12,     # 10 pM
    100e-12,    # 100 pM
    1e-9,       # 1 nM
    10e-9,      # 10 nM
    100e-9      # 100 nM
], dtype=float)

E2_FLOW_UL_MIN = 2.0
BUFFER_WASH_FLOWS_UL_MIN = [2.0, 4.0, 6.0, 8.0, 10.0]

INITIAL_BUFFER_MIN = 30.0
E2_PULSE_MIN = 30.0
WASH_MIN = 30.0


# ==============================================================================
# Working electrode and chamber geometry
# ==============================================================================

WE_DIAMETER_MM = 4.0
CHAMBER_DIAMETER_MM = 9.0
CHAMBER_HEIGHT_MM = 0.6

A_WE_MM2 = math.pi * (WE_DIAMETER_MM / 2.0) ** 2
A_WE_M2 = A_WE_MM2 * 1e-6

V_CH_UL = math.pi * (CHAMBER_DIAMETER_MM / 2.0) ** 2 * CHAMBER_HEIGHT_MM
V_CH_L = V_CH_UL * 1e-6
```

```
# ==============================================================================
#  Simulation parameters
# ==============================================================================

PARAMS = {
    # Binding parameters.
    "kon_M_inv_s": 5.0e5,       # apparent association rate, M^-1 s^-1
    "Kd_M": 5.0e-9,             # apparent dissociation constant, M
    "Gamma_pmol_cm2": 1.0,      # active aptamer coverage, pmol/cm^2

    # Mass-transfer parameter.
    "km_ref_um_s": 20.0,        # reference mass-transfer coefficient at Q_ref
    "Q_ref_uL_min": 2.0,        # reference flow for km scaling

    # Current mapping parameters.
    "I_unbound_nA": -230.0,     # unbound baseline current
    "DeltaI_nA": 80.0,          # full-scale binding-induced current change
    "drift_nA_per_min": 0.0,    # set to zero to isolate regeneration behavior

    # Initial conditions and numerical step.
    "theta0": 0.0,
    "Cb0_M": 0.0,
    "dt_s": 5.0
}


# ==============================================================================
# Helper functions
# ==============================================================================

def concentration_label(C_M):
    """Return a readable concentration label."""
    if C_M < 1e-9:
        return f"{C_M / 1e-12:.0f} pM"
    return f"{C_M / 1e-9:.0f} nM"


def flow_label(flow):
    """Return an Excel/Python-friendly flow-rate label."""
    return f"{int(flow) if float(flow).is_integer() else flow:g}uL_min"


def km_m_s(flow_uL_min, params):
    """
    Flow-dependent mass-transfer coefficient.
    Approximate laminar scaling:
        km(Q) = km_ref * (Q/Q_ref)^(1/3)
    """
    km_ref_m_s = params["km_ref_um_s"] * 1e-6
    return km_ref_m_s * (max(flow_uL_min, 1e-30) / params["Q_ref_uL_min"]) ** (1.0 / 3.0)


def calculate_Cs(theta, Cb_M, flow_uL_min, params):
    """
    Calculate interfacial E2 concentration Cs from the surface mass-transfer balance.

    1000*km*(Cb - Cs) =
    Gamma_T * [kon*Cs*(1 - theta) - koff*theta]

    The factor 1000 converts M = mol/L to mol/m^3 for compatibility with SI flux units.
    """
    kon = params["kon_M_inv_s"]
    Kd = params["Kd_M"]
    koff = kon * Kd

    Gamma_mol_m2 = params["Gamma_pmol_cm2"] * 1e-8
    km = km_m_s(flow_uL_min, params)

    theta = min(max(theta, 0.0), 1.0)
    Cb_M = max(Cb_M, 0.0)

    numerator = 1000.0 * km * Cb_M + Gamma_mol_m2 * koff * theta
    denominator = 1000.0 * km + Gamma_mol_m2 * kon * (1.0 - theta)

    return max(numerator / denominator, 0.0)
```

```
def derivatives(theta, Cb_M, Cin_M, flow_uL_min, params):
    """
    Coupled binding-transport model.

    Returns:
    dtheta/dt, dCb/dt, Cs
    """
    kon = params["kon_M_inv_s"]
    Kd = params["Kd_M"]
    koff = kon * Kd

    theta = min(max(theta, 0.0), 1.0)
    Cb_M = max(Cb_M, 0.0)

    Cs_M = calculate_Cs(theta, Cb_M, flow_uL_min, params)

    dtheta_dt = kon * Cs_M * (1.0 - theta) - koff * theta

    # Q/Vch in s^-1. Q and Vch are both in uL-based units here.
    q_over_v_s = (flow_uL_min / V_CH_UL) / 60.0

    # Surface capacity expressed as equivalent chamber concentration, M.
    Gamma_mol_m2 = params["Gamma_pmol_cm2"] * 1e-8
    surface_capacity_M = A_WE_M2 * Gamma_mol_m2 / V_CH_L

    dCb_dt = q_over_v_s * (Cin_M - Cb_M) - surface_capacity_M * dtheta_dt

    return dtheta_dt, dCb_dt, Cs_M


def rk4_step(theta, Cb_M, dt_s, Cin_M, flow_uL_min, params):
    """Fourth-order Runge-Kutta integration step."""
    def f(th, cb):
        dtheta, dCb, _ = derivatives(th, cb, Cin_M, flow_uL_min, params)
        return np.array([dtheta, dCb], dtype=float)

    y = np.array([theta, Cb_M], dtype=float)

    k1 = f(y[0], y[1])
    k2 = f(y[0] + 0.5 * dt_s * k1[0], y[1] + 0.5 * dt_s * k1[1])
    k3 = f(y[0] + 0.5 * dt_s * k2[0], y[1] + 0.5 * dt_s * k2[1])
    k4 = f(y[0] + dt_s * k3[0], y[1] + dt_s * k3[1])

    y_next = y + (dt_s / 6.0) * (k1 + 2.0*k2 + 2.0*k3 + k4)

    theta_next = float(np.clip(y_next[0], 0.0, 1.0))
    Cb_next = float(max(y_next[1], 0.0))

    return theta_next, Cb_next


def current_from_theta(theta, t_min, params):
    """Map fractional occupancy to simulated SWV peak current."""
    return (
        params["I_unbound_nA"]
        + params["DeltaI_nA"] * theta
        + params["drift_nA_per_min"] * t_min
    )


def simulate_one_wash_flow(wash_flow_uL_min, params):
    """
    Simulate the full cyclic sequence for one buffer-wash flow rate.
    """
    dt_s = params["dt_s"]
    theta = params["theta0"]
    Cb_M = params["Cb0_M"]
    t_min = 0.0

    rows = []

    def record(step_name, cycle, dose_label, Cin_M, flow_uL_min):
        _, _, Cs_M = derivatives(theta, Cb_M, Cin_M, flow_uL_min, params)
        rows.append({
            "wash_flow_uL_min": wash_flow_uL_min,
```

```
            "time_min": t_min,
            "current_nA": current_from_theta(theta, t_min, params),
            "theta": theta,
            "Cb_M": Cb_M,
            "Cs_M": Cs_M,
            "Cin_M": Cin_M,
            "applied_flow_uL_min": flow_uL_min,
            "step_name": step_name,
            "cycle": cycle,
            "dose_label": dose_label
        })

    def advance(duration_min, Cin_M, flow_uL_min, step_name, cycle, dose_label):
        nonlocal theta, Cb_M, t_min
        n_steps = int(round(duration_min * 60.0 / dt_s))

        for i in range(n_steps + 1):
            if not rows or i > 0:
                record(step_name, cycle, dose_label, Cin_M, flow_uL_min)

            if i < n_steps:
                theta, Cb_M = rk4_step(theta, Cb_M, dt_s, Cin_M, flow_uL_min, params)
                t_min += dt_s / 60.0

    # Initial buffer segment
    advance(INITIAL_BUFFER_MIN, 0.0, wash_flow_uL_min, "initial_buffer", 0, "buffer")

    # Sequential E2/wash cycles
    for cycle, C_M in enumerate(E2_CONCENTRATIONS_M, start=1):
        dose = concentration_label(C_M)
        advance(E2_PULSE_MIN, float(C_M), E2_FLOW_UL_MIN, "E2", cycle, dose)
        advance(WASH_MIN, 0.0, wash_flow_uL_min, "wash", cycle, "buffer")

    return rows


def make_wide_current_csv(long_rows, csv_path):
    """Create a wide-format CSV with one current column per wash-flow rate."""
    wash_flows = sorted({r["wash_flow_uL_min"] for r in long_rows})
    times = sorted({r["time_min"] for r in long_rows})

    by_time_flow = {(r["time_min"], r["wash_flow_uL_min"]): r for r in long_rows}

    # Use the first wash-flow condition for shared protocol metadata.
    first_flow = wash_flows[0]
    metadata_by_time = {}
    for r in long_rows:
        if r["wash_flow_uL_min"] == first_flow:
            metadata_by_time[r["time_min"]] = {
                "time_min": r["time_min"],
                "step_name": r["step_name"],
                "cycle": r["cycle"],
                "dose_label": r["dose_label"],
                "Cin_M": r["Cin_M"]
            }

    headers = ["time_min", "step_name", "cycle", "dose_label", "Cin_M"]
    headers += [f"current_wash_{flow_label(f)}_nA" for f in wash_flows]

    wide_rows = []
    for t in times:
        row = dict(metadata_by_time[t])
        for f in wash_flows:
            row[f"current_wash_{flow_label(f)}_nA"] = by_time_flow[(t, f)]["current_nA"]
        wide_rows.append(row)

    with Path(csv_path).open("w", newline="", encoding="utf-8") as f:
        writer = csv.DictWriter(f, fieldnames=headers)
        writer.writeheader()
        writer.writerows(wide_rows)


def plot_figure8a(long_rows, png_path, pdf_path):
    """Generate Figure 8a-style cyclic current plot."""
    fig, ax = plt.subplots(figsize=(11.5, 5.5))
```

```
    for flow in BUFFER_WASH_FLOWS_UL_MIN:
        sub = [r for r in long_rows if r["wash_flow_uL_min"] == flow]
        ax.plot(
            [r["time_min"] for r in sub],
            [r["current_nA"] for r in sub],
            linewidth=2.0,
            label=f"Wash {flow:g} uL/min"
        )

    # Add vertical boundaries and dose labels using the 2 uL/min trace.
    ref = [r for r in long_rows if r["wash_flow_uL_min"] == BUFFER_WASH_FLOWS_UL_MIN[0]]

    # Find protocol segments.
    segments = []
    current_key = None
    start_t = None
    last_t = None
    for r in ref:
        key = (r["step_name"], r["cycle"], r["dose_label"])
        t = r["time_min"]
        if current_key is None:
            current_key = key
            start_t = t
        elif key != current_key:
            segments.append((current_key[0], current_key[1], current_key[2], start_t, last_t))
            current_key = key
            start_t = t
        last_t = t
    segments.append((current_key[0], current_key[1], current_key[2], start_t, last_t))

    for _, _, _, t0, _ in segments:
        ax.axvline(t0, linewidth=0.8, alpha=0.18)

    ymin, ymax = ax.get_ylim()
    y_text = ymax - 0.05 * (ymax - ymin)
    for step_name, cycle, dose_label, t0, t1 in segments:
        if step_name == "E2":
            ax.text(
                0.5 * (t0 + t1),
                y_text,
                dose_label,
                ha="center",
                va="top",
                rotation=90,
                fontsize=8
            )

    ax.set_title("Simulated cyclic estradiol EAB operation vs. buffer-wash flow rate")
    ax.set_xlabel("Time (min)")
    ax.set_ylabel("Simulated SWV peak current (nA)")
    ax.grid(True, alpha=0.25)
    ax.legend(ncol=2, fontsize=9)
    fig.tight_layout()

    fig.savefig(png_path, dpi=300, bbox_inches="tight")
    fig.savefig(pdf_path, bbox_inches="tight")
    plt.show()


def main():
    output_dir = Path("figure8a_outputs")
    output_dir.mkdir(exist_ok=True)

    long_rows = []
    for wash_flow in BUFFER_WASH_FLOWS_UL_MIN:
        long_rows.extend(simulate_one_wash_flow(wash_flow, PARAMS))

    # Save long-format trace CSV.
    long_csv = output_dir / "Figure8a_time_traces_long.csv"
    with long_csv.open("w", newline="", encoding="utf-8") as f:
        writer = csv.DictWriter(f, fieldnames=list(long_rows[0].keys()))
        writer.writeheader()
        writer.writerows(long_rows)

    # Save wide-format current CSV.
    wide_csv = output_dir / "Figure8a_current_wide_by_wash_flow.csv"
```

```
    make_wide_current_csv(long_rows, wide_csv)

    # Plot Figure 8a.
    png_path = output_dir / "Figure8a_simulated_cyclic_current.png"
    pdf_path = output_dir / "Figure8a_simulated_cyclic_current.pdf"
    plot_figure8a(long_rows, png_path, pdf_path)

    # Print key values.
    koff = PARAMS["kon_M_inv_s"] * PARAMS["Kd_M"]
    t_half_min = math.log(2.0) / koff / 60.0

    print("Simulation complete.")
    print(f"Output folder: {output_dir.resolve()}")
    print(f"Working electrode area: {A_WE_MM2:.4f} mm^2")
    print(f"Chamber volume: {V_CH_UL:.4f} uL")
    print(f"koff = {koff:.3e} s^-1")
    print(f"t1/2,off = {t_half_min:.2f} min")


if __name__ == "__main__":
    main()
```

### 9. Developed code for numerical modeling of geometric washout (Figure 7b)

```
"""
Figure 8b simulation code:
Geometric chamber washout analysis for flow-programmed regeneration.

This script calculates the residual bulk E2 percentage remaining in the chamber
after buffer washing as a function of wash-flow rate and wash duration.

Model:
    residual fraction = exp(-(Q * t_w) / V_ch)

Device geometry:
    chamber diameter = 9 mm
    chamber height = 0.6 mm
    V_ch = 38.17 uL

Outputs:
    figure8b_outputs_linear/Figure8b_geometric_washout_percent_linear.png
    figure8b_outputs_linear/Figure8b_geometric_washout_percent_linear.pdf
    figure8b_outputs_linear/Figure8b_geometric_washout_summary.csv
    figure8b_outputs_linear/Figure8b_geometric_washout_wide.csv
"""

from pathlib import Path
import csv
import math
import matplotlib.pyplot as plt


# ============================================================================
# Experimental parameters
# ============================================================================

CHAMBER_DIAMETER_MM = 9.0
CHAMBER_HEIGHT_MM = 0.6

WASH_FLOWS_UL_MIN = [2, 4, 6, 8, 10]
WASH_DURATIONS_MIN = [5, 10, 15, 20, 25, 30, 35, 40, 45, 50, 55, 60]


# ============================================================================
# Geometry of chamber
# ============================================================================

V_CH_UL = math.pi * (CHAMBER_DIAMETER_MM / 2.0) ** 2 * CHAMBER_HEIGHT_MM


# ============================================================================
# Calculations
# ============================================================================

def calculate_washout_rows():
    rows = []

    for wash_time_min in WASH_DURATIONS_MIN:
        for flow_uL_min in WASH_FLOWS_UL_MIN:
            chamber_volumes_exchanged = flow_uL_min * wash_time_min / V_CH_UL
            residual_fraction = math.exp(-chamber_volumes_exchanged)

            rows.append({
                "wash_flow_uL_min": flow_uL_min,
                "wash_duration_min": wash_time_min,
                "Vch_uL": V_CH_UL,
                "chamber_volumes_exchanged": chamber_volumes_exchanged,
```

```
            "ideal_residual_fraction": residual_fraction,
            "ideal_residual_percent": 100.0 * residual_fraction,
        })

    return rows


def save_long_csv(rows, output_path):
    fieldnames = [
        "wash_flow_uL_min",
        "wash_duration_min",
        "Vch_uL",
        "chamber_volumes_exchanged",
        "ideal_residual_fraction",
        "ideal_residual_percent",
    ]

    with Path(output_path).open("w", newline="", encoding="utf-8") as f:
        writer = csv.DictWriter(f, fieldnames=fieldnames)
        writer.writeheader()
        writer.writerows(rows)


def save_wide_csv(rows, output_path):
    """
    Wide CSV format:
    one row per wash-flow rate and one residual-percent column per wash duration.
    """
    by_flow_time = {
        (row["wash_flow_uL_min"], row["wash_duration_min"]): row
        for row in rows
    }

    fieldnames = ["wash_flow_uL_min"]
    fieldnames += [f"residual_percent_{t}min" for t in WASH_DURATIONS_MIN]
    fieldnames += [f"residual_fraction_{t}min" for t in WASH_DURATIONS_MIN]

    wide_rows = []
    for flow in WASH_FLOWS_UL_MIN:
        out = {"wash_flow_uL_min": flow}
        for t in WASH_DURATIONS_MIN:
            out[f"residual_percent_{t}min"] = by_flow_time[(flow, t)]["ideal_residual_percent"]
        for t in WASH_DURATIONS_MIN:
            out[f"residual_fraction_{t}min"] = by_flow_time[(flow, t)]["ideal_residual_fraction"]
        wide_rows.append(out)

    with Path(output_path).open("w", newline="", encoding="utf-8") as f:
        writer = csv.DictWriter(f, fieldnames=fieldnames)
        writer.writeheader()
        writer.writerows(wide_rows)


def plot_washout_linear(rows, png_path, pdf_path):
    """
    Plot Figure 8b with a LINEAR y-axis.
    The plotted y-value is ideal residual bulk E2 percentage.
    """
    plt.figure(figsize=(10.8, 6.8))

    for wash_time in WASH_DURATIONS_MIN:
        x = []
        y = []
        for row in rows:
            if row["wash_duration_min"] == wash_time:
                x.append(row["wash_flow_uL_min"])
                y.append(row["ideal_residual_percent"])

        plt.plot(
            x,
            y,
            marker="o",
            linewidth=1.8,
            label=f"{wash_time} min wash"
        )

    plt.title("Geometric chamber washout scaling")
```

```
    plt.xlabel("Buffer-wash flow rate (uL/min)")
    plt.ylabel("Ideal residual bulk E2 (%)")
    plt.ylim(0, 100)
    plt.grid(True, alpha=0.25)
    plt.legend(ncol=2, fontsize=9)
    plt.tight_layout()

    plt.savefig(png_path, dpi=300, bbox_inches="tight")
    plt.savefig(pdf_path, bbox_inches="tight")
    plt.show()


def main():
    output_dir = Path("figure8b_outputs_linear")
    output_dir.mkdir(exist_ok=True)

    rows = calculate_washout_rows()

    long_csv = output_dir / "Figure8b_geometric_washout_summary.csv"
    wide_csv = output_dir / "Figure8b_geometric_washout_wide.csv"
    png_path = output_dir / "Figure8b_geometric_washout_percent_linear.png"
    pdf_path = output_dir / "Figure8b_geometric_washout_percent_linear.pdf"

    save_long_csv(rows, long_csv)
    save_wide_csv(rows, wide_csv)
    plot_washout_linear(rows, png_path, pdf_path)

    print("Figure 8b linear-axis geometric washout analysis complete.")
    print(f"Output folder: {output_dir.resolve()}")
    print(f"Chamber volume used: {V_CH_UL:.4f} uL")
    print(f"Long CSV: {long_csv}")
    print(f"Wide CSV: {wide_csv}")
    print(f"PNG: {png_path}")
    print(f"PDF: {pdf_path}")


if __name__ == "__main__":
    main()
```

#### 10. Developed code for numerical modeling of regeneration efficiency (Figure 7c)

```
"""
Figure 8c simulation code:
Regeneration efficiency extracted from the full coupled binding-transport model.

Model components:
1) Chamber exchange:
   dCb/dt = (Q/Vch)(Cin - Cb) - (A_WE*Gamma_T/Vch)(dtheta/dt)

2) Interfacial transport:
   km(Cb - Cs) = Gamma_T [kon*Cs(1 - theta) - koff*theta]

3) Surface occupancy:
   dtheta/dt = kon*Cs(1 - theta) - koff*theta

4) Current mapping:
   I(t) = I_unbound + DeltaI*theta(t)

5) Regeneration efficiency:
   eta_regen = 1 - |I_wash_end - I_start| / |I_E2_end - I_start|

Protocol:
- E2 sequence: 10 pM, 100 pM, 1 nM, 10 nM, 100 nM
- E2 pulse: 30 min at 2 uL/min
- Wash step: 30 min at 2, 4, 6, 8, or 10 uL/min
- Initial buffer: 30 min
"""

from pathlib import Path
import csv
import math
import numpy as np
import matplotlib.pyplot as plt


# ============================================================================
# Experiemntal Protocol
# ============================================================================

E2_CONCENTRATIONS_M = np.array([
    10e-12,     # 10 pM
    100e-12,    # 100 pM
    1e-9,       # 1 nM
    10e-9,      # 10 nM
    100e-9      # 100 nM
], dtype=float)

E2_FLOW_UL_MIN = 2.0
WASH_FLOWS_UL_MIN = [2.0, 4.0, 6.0, 8.0, 10.0]

INITIAL_BUFFER_MIN = 30.0
E2_PULSE_MIN = 30.0
WASH_MIN = 30.0


# ============================================================================
# Device geometry of working electrode and chamber
# ============================================================================

WE_DIAMETER_MM = 4.0
CHAMBER_DIAMETER_MM = 9.0
CHAMBER_HEIGHT_MM = 0.6
```

```
A_WE_MM2 = math.pi * (WE_DIAMETER_MM / 2.0) ** 2
A_WE_M2 = A_WE_MM2 * 1e-6

V_CH_UL = math.pi * (CHAMBER_DIAMETER_MM / 2.0) ** 2 * CHAMBER_HEIGHT_MM
V_CH_L = V_CH_UL * 1e-6


# ==========================================================================
#  Simulation parameters
# ==========================================================================

PARAMS = {
    # Representative apparent kinetic parameters for simulation/design-space analysis.
    "kon_M_inv_s": 5.0e5,
    "Kd_M": 5.0e-9,
    "Gamma_pmol_cm2": 1.0,

    # Mass-transfer coefficient at reference flow.
    "km_ref_um_s": 20.0,
    "Q_ref_uL_min": 2.0,

    # Current mapping.
    "I_unbound_nA": -230.0,
    "DeltaI_nA": 80.0,
    "drift_nA_per_min": 0.0,

    # Initial conditions and numerical step.
    "theta0": 0.0,
    "Cb0_M": 0.0,
    "dt_s": 5.0
}


# ==========================================================================
# Helper functions
# ==========================================================================

def concentration_label(C_M):
    if C_M < 1e-9:
        return f"{C_M / 1e-12:.0f} pM"
    return f"{C_M / 1e-9:.0f} nM"


def flow_label(flow):
    return f"{int(flow) if float(flow).is_integer() else flow:g}uL_min"


def km_m_s(flow_uL_min, params):
    return (
        params["km_ref_um_s"] * 1e-6
        * (max(flow_uL_min, 1e-30) / params["Q_ref_uL_min"]) ** (1.0 / 3.0)
    )


def calculate_Cs(theta, Cb_M, flow_uL_min, params):
    """
    Calculate interfacial E2 concentration Cs from the surface mass-transfer balance.

    1000*km*(Cb - Cs) =
    Gamma_T * [kon*Cs*(1 - theta) - koff*theta]
    """
    kon = params["kon_M_inv_s"]
    Kd = params["Kd_M"]
    koff = kon * Kd

    Gamma_mol_m2 = params["Gamma_pmol_cm2"] * 1e-8
    km = km_m_s(flow_uL_min, params)

    theta = min(max(theta, 0.0), 1.0)
    Cb_M = max(Cb_M, 0.0)

    numerator = 1000.0 * km * Cb_M + Gamma_mol_m2 * koff * theta
    denominator = 1000.0 * km + Gamma_mol_m2 * kon * (1.0 - theta)

    return max(numerator / denominator, 0.0)
```

```
def derivatives(theta, Cb_M, Cin_M, flow_uL_min, params):
    """
    Coupled chamber exchange and surface occupancy model.

    Returns:
        dtheta/dt, dCb/dt, Cs
    """
    kon = params["kon_M_inv_s"]
    Kd = params["Kd_M"]
    koff = kon * Kd

    theta = min(max(theta, 0.0), 1.0)
    Cb_M = max(Cb_M, 0.0)

    Cs_M = calculate_Cs(theta, Cb_M, flow_uL_min, params)

    dtheta_dt = kon * Cs_M * (1.0 - theta) - koff * theta

    q_over_v_s = (flow_uL_min / V_CH_UL) / 60.0

    Gamma_mol_m2 = params["Gamma_pmol_cm2"] * 1e-8
    surface_capacity_M = A_WE_M2 * Gamma_mol_m2 / V_CH_L

    dCb_dt = q_over_v_s * (Cin_M - Cb_M) - surface_capacity_M * dtheta_dt

    return dtheta_dt, dCb_dt, Cs_M


def rk4_step(theta, Cb_M, dt_s, Cin_M, flow_uL_min, params):
    """Fourth-order Runge-Kutta integration step."""
    def f(th, cb):
        dtheta, dCb, _ = derivatives(th, cb, Cin_M, flow_uL_min, params)
        return np.array([dtheta, dCb], dtype=float)

    y = np.array([theta, Cb_M], dtype=float)

    k1 = f(y[0], y[1])
    k2 = f(y[0] + 0.5 * dt_s * k1[0], y[1] + 0.5 * dt_s * k1[1])
    k3 = f(y[0] + 0.5 * dt_s * k2[0], y[1] + 0.5 * dt_s * k2[1])
    k4 = f(y[0] + dt_s * k3[0], y[1] + dt_s * k3[1])

    y_next = y + (dt_s / 6.0) * (k1 + 2.0*k2 + 2.0*k3 + k4)

    theta_next = float(np.clip(y_next[0], 0.0, 1.0))
    Cb_next = float(max(y_next[1], 0.0))

    return theta_next, Cb_next


def current_from_theta(theta, t_min, params):
    return (
        params["I_unbound_nA"]
        + params["DeltaI_nA"] * theta
        + params["drift_nA_per_min"] * t_min
    )


def simulate_one_wash_flow(wash_flow_uL_min, params):
    """
    Simulate the cyclic sequence for one wash-flow rate and calculate
    regeneration efficiency after each pulse.
    """
    dt_s = params["dt_s"]
    theta = params["theta0"]
    Cb_M = params["Cb0_M"]
    t_min = 0.0

    trace_rows = []
    metrics_rows = []

    def record(step_name, cycle, dose_label, Cin_M, flow_uL_min):
        _, _, Cs_M = derivatives(theta, Cb_M, Cin_M, flow_uL_min, params)
        trace_rows.append({
            "wash_flow_uL_min": wash_flow_uL_min,
            "time_min": t_min,
```

```
        "step_name": step_name,
        "cycle": cycle,
        "dose_label": dose_label,
        "Cin_M": Cin_M,
        "applied_flow_uL_min": flow_uL_min,
        "theta": theta,
        "Cb_M": Cb_M,
        "Cs_M": Cs_M,
        "current_nA": current_from_theta(theta, t_min, params),
    })

def advance(duration_min, Cin_M, flow_uL_min, step_name, cycle, dose_label):
    nonlocal theta, Cb_M, t_min
    n_steps = int(round(duration_min * 60.0 / dt_s))

    for i in range(n_steps + 1):
        if not trace_rows or i > 0:
            record(step_name, cycle, dose_label, Cin_M, flow_uL_min)

        if i < n_steps:
            theta, Cb_M = rk4_step(theta, Cb_M, dt_s, Cin_M, flow_uL_min, params)
            t_min += dt_s / 60.0

# Initial buffer
advance(INITIAL_BUFFER_MIN, 0.0, wash_flow_uL_min, "initial_buffer", 0, "buffer")

# E2/wash cycles
for cycle_idx, C_M in enumerate(E2_CONCENTRATIONS_M, start=1):
    dose = concentration_label(C_M)

    I_start = current_from_theta(theta, t_min, params)
    theta_start = theta
    Cb_start = Cb_M
    _, _, Cs_start = derivatives(theta, Cb_M, float(C_M), E2_FLOW_UL_MIN, params)
    t_start = t_min

    # E2 pulse
    advance(E2_PULSE_MIN, float(C_M), E2_FLOW_UL_MIN, "E2", cycle_idx, dose)
    I_E2_end = current_from_theta(theta, t_min, params)
    theta_E2_end = theta
    Cb_E2_end = Cb_M
    _, _, Cs_E2_end = derivatives(theta, Cb_M, float(C_M), E2_FLOW_UL_MIN, params)
    t_E2_end = t_min

    # Wash
    advance(WASH_MIN, 0.0, wash_flow_uL_min, "wash", cycle_idx, "buffer")
    I_wash_end = current_from_theta(theta, t_min, params)
    theta_wash_end = theta
    Cb_wash_end = Cb_M
    _, _, Cs_wash_end = derivatives(theta, Cb_M, 0.0, wash_flow_uL_min, params)
    t_wash_end = t_min

    response = abs(I_E2_end - I_start)
    residual = abs(I_wash_end - I_start)
    memory_fraction = residual / response if response > 0 else np.nan
    regeneration_efficiency = 1.0 - memory_fraction if response > 0 else np.nan

    metrics_rows.append({
        "wash_flow_uL_min": wash_flow_uL_min,
        "cycle": cycle_idx,
        "E2_label": dose,
        "E2_M": float(C_M),
        "t_start_min": t_start,
        "t_E2_end_min": t_E2_end,
        "t_wash_end_min": t_wash_end,
        "I_start_nA": I_start,
        "I_E2_end_nA": I_E2_end,
        "I_wash_end_nA": I_wash_end,
        "response_nA": response,
        "residual_after_wash_nA": residual,
        "memory_fraction": memory_fraction,
        "regeneration_efficiency": regeneration_efficiency,
        "regeneration_efficiency_percent": 100.0 * regeneration_efficiency,
        "theta_start": theta_start,
        "theta_E2_end": theta_E2_end,
        "theta_wash_end": theta_wash_end,
```

```
            "Cb_start_M": Cb_start,
            "Cb_E2_end_M": Cb_E2_end,
            "Cb_wash_end_M": Cb_wash_end,
            "Cs_start_M": Cs_start,
            "Cs_E2_end_M": Cs_E2_end,
            "Cs_wash_end_M": Cs_wash_end,
        })

    return trace_rows, metrics_rows


def save_wide_csv(metrics_rows, output_path):
    dose_labels = [concentration_label(c) for c in E2_CONCENTRATIONS_M]
    by_dose_flow = {
        (row["E2_label"], row["wash_flow_uL_min"]): row
        for row in metrics_rows
    }

    fieldnames = ["E2_label", "E2_M"]
    fieldnames += [f"regen_efficiency_percent_wash_{flow_label(flow)}" for flow in WASH_FLOWS_UL_MIN]
    fieldnames += [f"memory_fraction_wash_{flow_label(flow)}" for flow in WASH_FLOWS_UL_MIN]

    wide_rows = []
    for C_M, dose in zip(E2_CONCENTRATIONS_M, dose_labels):
        out = {
            "E2_label": dose,
            "E2_M": float(C_M),
        }

        for flow in WASH_FLOWS_UL_MIN:
            out[f"regen_efficiency_percent_wash_{flow_label(flow)}"] = (
                by_dose_flow[(dose, flow)]["regeneration_efficiency_percent"]
            )

        for flow in WASH_FLOWS_UL_MIN:
            out[f"memory_fraction_wash_{flow_label(flow)}"] = (
                by_dose_flow[(dose, flow)]["memory_fraction"]
            )

        wide_rows.append(out)

    with Path(output_path).open("w", newline="", encoding="utf-8") as f:
        writer = csv.DictWriter(f, fieldnames=fieldnames)
        writer.writeheader()
        writer.writerows(wide_rows)


def plot_figure8c(metrics_rows, png_path, pdf_path):
    dose_labels = [concentration_label(c) for c in E2_CONCENTRATIONS_M]
    by_dose_flow = {
        (row["E2_label"], row["wash_flow_uL_min"]): row
        for row in metrics_rows
    }

    fig, ax = plt.subplots(figsize=(8.7, 5.2))
    x = np.arange(len(dose_labels))

    for flow in WASH_FLOWS_UL_MIN:
        y = [
            by_dose_flow[(dose, flow)]["regeneration_efficiency"]
            for dose in dose_labels
        ]

        ax.plot(
            x,
            y,
            marker="o",
            linewidth=2.0,
            label=f"Wash {flow:g} uL/min"
        )

    ax.set_title("Full coupled-model regeneration efficiency")
    ax.set_xlabel("Estradiol concentration per cycle")
    ax.set_ylabel("Regeneration efficiency")
    ax.set_xticks(x)
    ax.set_xticklabels(dose_labels)
```

```
    ax.set_ylim(0.0, 1.05)
    ax.grid(True, alpha=0.25)
    ax.legend(ncol=2, fontsize=9)

    fig.tight_layout()
    fig.savefig(png_path, dpi=300, bbox_inches="tight")
    fig.savefig(pdf_path, bbox_inches="tight")
    plt.show()


def main():
    output_dir = Path("figure8c_outputs")
    output_dir.mkdir(exist_ok=True)

    all_trace_rows = []
    all_metrics_rows = []

    for wash_flow in WASH_FLOWS_UL_MIN:
        trace_rows, metrics_rows = simulate_one_wash_flow(wash_flow, PARAMS)
        all_trace_rows.extend(trace_rows)
        all_metrics_rows.extend(metrics_rows)

    # Save metrics long CSV.
    metrics_csv = output_dir / "Figure8c_regeneration_metrics_long.csv"
    with metrics_csv.open("w", newline="", encoding="utf-8") as f:
        writer = csv.DictWriter(f, fieldnames=list(all_metrics_rows[0].keys()))
        writer.writeheader()
        writer.writerows(all_metrics_rows)

    # Save time traces CSV.
    traces_csv = output_dir / "Figure8c_time_traces_full_model.csv"
    with traces_csv.open("w", newline="", encoding="utf-8") as f:
        writer = csv.DictWriter(f, fieldnames=list(all_trace_rows[0].keys()))
        writer.writeheader()
        writer.writerows(all_trace_rows)

    # Save wide CSV.
    wide_csv = output_dir / "Figure8c_regeneration_efficiency_wide.csv"
    save_wide_csv(all_metrics_rows, wide_csv)

    # Plot.
    png_path = output_dir / "Figure8c_full_coupled_regeneration_efficiency.png"
    pdf_path = output_dir / "Figure8c_full_coupled_regeneration_efficiency.pdf"
    plot_figure8c(all_metrics_rows, png_path, pdf_path)

    # Print key values.
    koff = PARAMS["kon_M_inv_s"] * PARAMS["Kd_M"]
    t_half_min = math.log(2.0) / koff / 60.0

    print("Figure 8c full coupled-model simulation complete.")
    print(f"Output folder: {output_dir.resolve()}")
    print(f"Working electrode area: {A_WE_MM2:.4f} mm^2")
    print(f"Chamber volume: {V_CH_UL:.4f} uL")
    print(f"koff = {koff:.3e} s^-1")
    print(f"t1/2,off = {t_half_min:.2f} min")
    print(f"Metrics CSV: {metrics_csv}")
    print(f"Wide CSV: {wide_csv}")
    print(f"Trace CSV: {traces_csv}")
    print(f"PNG: {png_path}")
    print(f"PDF: {pdf_path}")


if __name__ == "__main__":
    main()
```

### 11. Developed code for numerical modeling of $K_d$ affect on regeneration efficiency (Figure 7d, e, and f)

```
"""
Figure 7d–f combined heatmap code:
Regeneration-efficiency design maps for three representative apparent Kd values.

This code generates one combined figure containing three heatmaps:
    Figure 7d: Kd = 20 nM
    Figure 7e: Kd = 5 nM
    Figure 7f: Kd = 1 nM

Metric plotted:
    Minimum regeneration efficiency (%) across the full E2 sequence.

Sequence:
    10 pM, 100 pM, 1 nM, 10 nM, 100 nM

Protocol:
    E2 pulse = 30 min at 2 uL/min
    Buffer-wash flow = 2, 4, 6, 8, 10 uL/min
    Buffer-wash duration = 5, 10, 15, 20, 25, 30, 35, 40, 45, 50, 55, 60 min

Model components:
    1) Chamber exchange
    2) Interfacial transport
    3) Aptamer occupancy
    4) Rebinding/dissociation
    5) Current mapping
    6) Regeneration-efficiency extraction

Outputs:
    figure8def_outputs/Figure8def_combined_Kd_heatmaps_with_numbers.png
    figure8def_outputs/Figure8def_combined_Kd_heatmaps_with_numbers.pdf
    figure8def_outputs/Figure8def_combined_Kd_heatmaps_without_numbers.png
    figure8def_outputs/Figure8def_combined_Kd_heatmaps_without_numbers.pdf
    figure8def_outputs/Figure8def_Kd_heatmap_data_long.csv
    figure8def_outputs/Figure8def_Kd_heatmap_data_wide.csv
"""

from pathlib import Path
import csv
import math
import numpy as np
import matplotlib.pyplot as plt


# ============================================================================
# Experimental protocol and device geometry
# ============================================================================

E2_CONCENTRATIONS_M = np.array([
    10e-12,     # 10 pM
    100e-12,    # 100 pM
    1e-9,       # 1 nM
    10e-9,      # 10 nM
    100e-9      # 100 nM
], dtype=float)

E2_FLOW_UL_MIN = 2.0
INITIAL_BUFFER_MIN = 30.0
E2_PULSE_MIN = 30.0

WASH_FLOWS_UL_MIN = [2, 4, 6, 8, 10]
WASH_DURATIONS_MIN = [5, 10, 15, 20, 25, 30, 35, 40, 45, 50, 55, 60]
```

```
WE_DIAMETER_MM = 4.0
CHAMBER_DIAMETER_MM = 9.0
CHAMBER_HEIGHT_MM = 0.6

A_WE_MM2 = math.pi * (WE_DIAMETER_MM / 2.0) ** 2
A_WE_M2 = A_WE_MM2 * 1e-6

V_CH_UL = math.pi * (CHAMBER_DIAMETER_MM / 2.0) ** 2 * CHAMBER_HEIGHT_MM
V_CH_L = V_CH_UL * 1e-6


# ============================================================================
# Simulation parameters
# ============================================================================

BASE_PARAMS = {
    "kon_app_M_inv_s": 5.0e5,      # representative apparent association rate
    "Gamma_pmol_cm2": 1.0,         # representative active aptamer surface density
    "km_ref_um_s": 20.0,           # reference mass-transfer coefficient at Q_ref
    "Q_ref_uL_min": 2.0,           # reference flow rate for km scaling
    "I_unbound_nA": -230.0,        # unbound-state current
    "DeltaI_nA": 80.0,             # full-scale binding-induced current change
    "drift_nA_per_min": 0.0,       # set to zero to isolate regeneration physics
    "theta0": 0.0,
    "Cb0_M": 0.0,
    "dt_s": 5.0
}

KD_CASES = [
    {
        "panel": "d",
        "case_name": "Lower affinity",
        "Kd_nM": 20.0,
        "Kd_M": 20e-9
    },
    {
        "panel": "e",
        "case_name": "Intermediate affinity",
        "Kd_nM": 5.0,
        "Kd_M": 5e-9
    },
    {
        "panel": "f",
        "case_name": "Higher affinity",
        "Kd_nM": 1.0,
        "Kd_M": 1e-9
    }
]


# ============================================================================
# Model functions
# ============================================================================

def km_m_s(flow_uL_min, params):
    """
    Approximate flow-dependent mass-transfer coefficient:
        km(Q) = km_ref * (Q/Q_ref)^(1/3)
    """
    return (
        params["km_ref_um_s"] * 1e-6
        * (max(flow_uL_min, 1e-30) / params["Q_ref_uL_min"]) ** (1.0 / 3.0)
    )


def calculate_Cs(theta, Cb_M, flow_uL_min, params):
    """
    Interfacial mass-transfer balance:
        1000*km*(Cb - Cs) =
        Gamma_T * [kon*Cs*(1 - theta) - koff*theta]

    Rearranged:
        Cs = (1000*km*Cb + Gamma_T*koff*theta) /
             (1000*km + Gamma_T*kon*(1 - theta))
    """
```

```
    kon = params["kon_app_M_inv_s"]
    koff = kon * params["Kd_M"]
    Gamma_mol_m2 = params["Gamma_pmol_cm2"] * 1e-8
    km = km_m_s(flow_uL_min, params)

    theta = min(max(theta, 0.0), 1.0)
    Cb_M = max(Cb_M, 0.0)

    numerator = 1000.0 * km * Cb_M + Gamma_mol_m2 * koff * theta
    denominator = 1000.0 * km + Gamma_mol_m2 * kon * (1.0 - theta)

    return max(numerator / denominator, 0.0)


def derivatives(theta, Cb_M, Cin_M, flow_uL_min, params):
    """
    Full coupled binding-transport model.

    Surface occupancy:
        dtheta/dt = kon*Cs*(1 - theta) - koff*theta

    Chamber mass balance:
        dCb/dt = (Q/Vch)*(Cin - Cb) - (A_WE*Gamma_T/Vch)*(dtheta/dt)
    """
    kon = params["kon_app_M_inv_s"]
    koff = kon * params["Kd_M"]

    theta = min(max(theta, 0.0), 1.0)
    Cb_M = max(Cb_M, 0.0)

    Cs_M = calculate_Cs(theta, Cb_M, flow_uL_min, params)

    dtheta_dt = kon * Cs_M * (1.0 - theta) - koff * theta

    q_over_v_s = (flow_uL_min / V_CH_UL) / 60.0

    Gamma_mol_m2 = params["Gamma_pmol_cm2"] * 1e-8
    surface_capacity_M = A_WE_M2 * Gamma_mol_m2 / V_CH_L

    dCb_dt = q_over_v_s * (Cin_M - Cb_M) - surface_capacity_M * dtheta_dt

    return dtheta_dt, dCb_dt, Cs_M


def rk4_step(theta, Cb_M, dt_s, Cin_M, flow_uL_min, params):
    """Fourth-order Runge-Kutta integration step."""
    def f(th, cb):
        dtheta, dCb, _ = derivatives(th, cb, Cin_M, flow_uL_min, params)
        return np.array([dtheta, dCb], dtype=float)

    y = np.array([theta, Cb_M], dtype=float)

    k1 = f(y[0], y[1])
    k2 = f(y[0] + 0.5 * dt_s * k1[0], y[1] + 0.5 * dt_s * k1[1])
    k3 = f(y[0] + 0.5 * dt_s * k2[0], y[1] + 0.5 * dt_s * k2[1])
    k4 = f(y[0] + dt_s * k3[0], y[1] + dt_s * k3[1])

    y_next = y + (dt_s / 6.0) * (k1 + 2.0*k2 + 2.0*k3 + k4)

    theta_next = float(np.clip(y_next[0], 0.0, 1.0))
    Cb_next = float(max(y_next[1], 0.0))

    return theta_next, Cb_next


def current_from_theta(theta, t_min, params):
    """Current mapping: I(t) = I_unbound + DeltaI*theta + drift*t."""
    return (
        params["I_unbound_nA"]
        + params["DeltaI_nA"] * theta
        + params["drift_nA_per_min"] * t_min
    )


def advance_segment(theta, Cb_M, duration_min, Cin_M, flow_uL_min, params, t_start_min):
    """
```

```
    Advance theta and Cb through one protocol segment.
    Returns updated theta, Cb, segment-start current, segment-end current, and end time.
    """
    dt_s = params["dt_s"]
    n_steps = int(round(duration_min * 60.0 / dt_s))

    I_start = current_from_theta(theta, t_start_min, params)

    for _ in range(n_steps):
        theta, Cb_M = rk4_step(theta, Cb_M, dt_s, Cin_M, flow_uL_min, params)

    t_end_min = t_start_min + duration_min
    I_end = current_from_theta(theta, t_end_min, params)

    return theta, Cb_M, I_start, I_end, t_end_min


def simulate_sequence_for_condition(wash_flow_uL_min, wash_duration_min, params):
    """
    Simulate the full 10 pM to 100 nM E2 sequence for one wash-flow/wash-duration
    condition and return regeneration metrics.
    """
    theta = params["theta0"]
    Cb_M = params["Cb0_M"]
    t_min = 0.0

    # Initial buffer equilibration
    theta, Cb_M, _, _, t_min = advance_segment(
        theta=theta,
        Cb_M=Cb_M,
        duration_min=INITIAL_BUFFER_MIN,
        Cin_M=0.0,
        flow_uL_min=wash_flow_uL_min,
        params=params,
        t_start_min=t_min
    )

    regen_efficiencies = []
    memory_fractions = []

    for C_M in E2_CONCENTRATIONS_M:
        # Baseline before E2 pulse
        I_baseline = current_from_theta(theta, t_min, params)

        # E2 pulse at fixed E2 flow
        theta, Cb_M, _, I_E2_end, t_min = advance_segment(
            theta=theta,
            Cb_M=Cb_M,
            duration_min=E2_PULSE_MIN,
            Cin_M=float(C_M),
            flow_uL_min=E2_FLOW_UL_MIN,
            params=params,
            t_start_min=t_min
        )

        # Buffer wash at variable wash flow and duration
        theta, Cb_M, _, I_wash_end, t_min = advance_segment(
            theta=theta,
            Cb_M=Cb_M,
            duration_min=wash_duration_min,
            Cin_M=0.0,
            flow_uL_min=wash_flow_uL_min,
            params=params,
            t_start_min=t_min
        )

        response = abs(I_E2_end - I_baseline)
        residual = abs(I_wash_end - I_baseline)

        memory_fraction = residual / response if response > 0 else np.nan
        regen_eff = 100.0 * (1.0 - memory_fraction) if response > 0 else np.nan

        regen_efficiencies.append(regen_eff)
        memory_fractions.append(100.0 * memory_fraction if not np.isnan(memory_fraction) else np.nan)

    min_regen = float(np.nanmin(regen_efficiencies))
```

```
    min_regen_clipped = max(0.0, min_regen)

    return {
        "min_regeneration_efficiency_percent": min_regen,
        "min_regeneration_efficiency_percent_clipped": min_regen_clipped,
        "mean_regeneration_efficiency_percent": float(np.nanmean(regen_efficiencies)),
        "final_regeneration_efficiency_percent": float(regen_efficiencies[-1]),
        "max_memory_fraction_percent": float(np.nanmax(memory_fractions)),
        "mean_memory_fraction_percent": float(np.nanmean(memory_fractions))
    }


# ============================================================================
# Simulation and output functions
# ============================================================================

def run_all_heatmaps():
    all_rows = []
    heatmaps = {}

    for case in KD_CASES:
        params = dict(BASE_PARAMS)
        params["Kd_M"] = case["Kd_M"]

        heat = np.zeros((len(WASH_FLOWS_UL_MIN), len(WASH_DURATIONS_MIN)))

        for i, flow in enumerate(WASH_FLOWS_UL_MIN):
            for j, duration in enumerate(WASH_DURATIONS_MIN):
                metrics = simulate_sequence_for_condition(flow, duration, params)
                heat[i, j] = metrics["min_regeneration_efficiency_percent_clipped"]

                all_rows.append({
                    "panel": case["panel"],
                    "case_name": case["case_name"],
                    "Kd_nM": case["Kd_nM"],
                    "kon_app_M_inv_s": params["kon_app_M_inv_s"],
                    "koff_app_s_inv": params["kon_app_M_inv_s"] * params["Kd_M"],
                    "wash_flow_uL_min": flow,
                    "wash_duration_min": duration,
                    "sequence": "10 pM, 100 pM, 1 nM, 10 nM, 100 nM",
                    **metrics
                })

        heatmaps[case["panel"]] = {
            "case": case,
            "heat": heat
        }

    return heatmaps, all_rows


def save_long_csv(rows, output_path):
    fieldnames = list(rows[0].keys())

    with Path(output_path).open("w", newline="", encoding="utf-8") as f:
        writer = csv.DictWriter(f, fieldnames=fieldnames)
        writer.writeheader()
        writer.writerows(rows)


def save_wide_csv(rows, output_path):
    """
    Save heatmap values in wide format:
    one row per Kd/flow combination, columns = wash durations.
    """
    fieldnames = ["panel", "case_name", "Kd_nM", "wash_flow_uL_min"]
    fieldnames += [f"regen_eff_percent_{duration}min" for duration in WASH_DURATIONS_MIN]

    wide_rows = []

    for case in KD_CASES:
        for flow in WASH_FLOWS_UL_MIN:
            out = {
                "panel": case["panel"],
                "case_name": case["case_name"],
                "Kd_nM": case["Kd_nM"],
```

```
            "wash_flow_uL_min": flow
        }

        for duration in WASH_DURATIONS_MIN:
            match = [
                row for row in rows
                if row["panel"] == case["panel"]
                and row["wash_flow_uL_min"] == flow
                and row["wash_duration_min"] == duration
            ][0]
            out[f"regen_eff_percent_{duration}min"] = match["min_regeneration_efficiency_percent_clipped"]

        wide_rows.append(out)

    with Path(output_path).open("w", newline="", encoding="utf-8") as f:
        writer = csv.DictWriter(f, fieldnames=fieldnames)
        writer.writeheader()
        writer.writerows(wide_rows)


def plot_combined_heatmaps(heatmaps, output_png, output_pdf, annotate=True):
    """
    Generate one combined figure containing the three Kd heatmaps.
    """
    fig, axes = plt.subplots(1, 3, figsize=(16.0, 5.4), constrained_layout=True)

    panel_order = ["d", "e", "f"]

    for ax, panel in zip(axes, panel_order):
        heat = heatmaps[panel]["heat"]
        case = heatmaps[panel]["case"]

        im = ax.imshow(heat, aspect="auto", origin="lower", vmin=0, vmax=100)

        ax.set_title(
            f"({panel}) {case['case_name']}\nKd = {case['Kd_nM']:.0f} nM",
            fontsize=12
        )
        ax.set_xlabel("Wash duration (min)")
        ax.set_xticks(np.arange(len(WASH_DURATIONS_MIN)))
        ax.set_xticklabels(WASH_DURATIONS_MIN, rotation=45, ha="right")

        ax.set_yticks(np.arange(len(WASH_FLOWS_UL_MIN)))
        ax.set_yticklabels(WASH_FLOWS_UL_MIN)

        if panel == "d":
            ax.set_ylabel("Wash flow rate (uL/min)")
        else:
            ax.set_ylabel("")

        if annotate:
            for i in range(heat.shape[0]):
                for j in range(heat.shape[1]):
                    val = heat[i, j]
                    text_color = "white" if val < 55 else "black"
                    ax.text(
                        j,
                        i,
                        f"{val:.0f}%",
                        ha="center",
                        va="center",
                        fontsize=7.3,
                        color=text_color
                    )

    cbar = fig.colorbar(im, ax=axes, shrink=0.88)
    cbar.set_label("Minimum regeneration efficiency (%)")

    fig.suptitle(
        "Regeneration-efficiency maps for representative aptamer affinity regimes",
        fontsize=14
    )

    fig.savefig(output_png, dpi=300, bbox_inches="tight")
    fig.savefig(output_pdf, bbox_inches="tight")
    plt.show()
```

```
def main():
    output_dir = Path("figure8def_outputs")
    output_dir.mkdir(exist_ok=True)

    heatmaps, rows = run_all_heatmaps()

    long_csv = output_dir / "Figure8def_Kd_heatmap_data_long.csv"
    wide_csv = output_dir / "Figure8def_Kd_heatmap_data_wide.csv"

    save_long_csv(rows, long_csv)
    save_wide_csv(rows, wide_csv)

    # Combined figure with cell values.
    plot_combined_heatmaps(
        heatmaps,
        output_png=output_dir / "Figure8def_combined_Kd_heatmaps_with_numbers.png",
        output_pdf=output_dir / "Figure8def_combined_Kd_heatmaps_with_numbers.pdf",
        annotate=True
    )


    print("Figure 8d–f combined heatmap simulation complete.")
    print(f"Output folder: {output_dir.resolve()}")
    print(f"Chamber volume: {V_CH_UL:.4f} uL")
    print(f"Working electrode area: {A_WE_MM2:.4f} mm^2")
    print(f"Long CSV: {long_csv}")
    print(f"Wide CSV: {wide_csv}")

    for case in KD_CASES:
        koff = BASE_PARAMS["kon_app_M_inv_s"] * case["Kd_M"]
        half_life_min = math.log(2.0) / koff / 60.0
        print(
            f"Panel {case['panel']}: Kd = {case['Kd_nM']:.0f} nM, "
            f"koff = {koff:.3e} s^-1, "
            f"t1/2 = {half_life_min:.2f} min"
        )


if __name__ == "__main__":
    main()
```

**Table 1. Device geometry and flow protocol parameters.**

| Parameter | Symbol / code name | Value used | Unit | Use in model |
|---|---|---|---|---|
| **Chamber diameter** | d_ch; CHAMBER_DIAMETER_MM | 9.0 | mm | Defines chamber volume used in exchange and washout calculations |
| **Chamber height** | h; CHAMBER_HEIGHT_MM | 0.6 | mm | Defines chamber volume used in exchange and washout calculations |
| **Chamber volume** | V_ch; V_CH_UL | 38.17 | µL | Chamber exchange, residence time, ideal washout, and coupled transport model |
| **Working-electrode diameter** | d_WE; WE_DIAMETER_MM | 4.0 | mm | Defines active working-electrode area |
| **Working-electrode area** | A_WE; A_WE_MM2 | 12.57 | $mm^2$ | Surface uptake/release term and total aptamer capacity |
| **E2 concentration sequence** | C_in; E2_CONCENTRATIONS_M | 10 pM, 100 pM, 1 nM, 10 nM, 100 nM | M | Sequential target pulses for Figure 8a,c,d–f simulations |
| **E2 injection flow rate** | Q_E2; E2_FLOW_UL_MIN | 2.0 | µL $min^{-1}$ | Target delivery during E2 exposure steps |
| **Buffer-wash flow rates** | Q_wash; WASH_FLOWS_UL_MIN | 2, 4, 6, 8, 10 | µL $min^{-1}$ | Controls chamber exchange and regeneration during wash steps |
| **Initial buffer step** | INITIAL_BUFFER_MIN | 30 | min | Establishes the initial simulated baseline |
| **E2 pulse duration** | t_p; E2_PULSE_MIN | 30 | min | Association/exposure interval for each target dose |
| **Wash duration for Figure 8a,c** | t_w; WASH_MIN | 30 | min | Regeneration interval after each target pulse |
| **Wash durations for Figure 8b,d–f** | t_w; WASH_DURATIONS_MIN | 5, 10, 15, 20, 25, 30, 35, 40, 45, 50, 55, 60 | min | Flow-duration design-space mapping |

**Table 2. Binding, chamber-transport, surface-capacity, and interfacial-transport parameters.**

| Parameter | Symbol / code name | Value used | Unit | Use in model |
|---|---|---|---|---|
| **Bulk chamber E2 concentration** | C_b(t) | Simulated state variable | M | Bulk concentration in the chamber mass balance |
| **Inlet E2 concentration** | C_in(t) | 0 during wash; E2 dose during pulse | M | Driving concentration in chamber exchange |
| **Chamber exchange rate** | Q/V_ch | Derived | $s^{-1}$ | Controls chamber clearance and residence time |
| **Ideal residual fraction** | C_b(t_w)/C_b(0) | Derived | dimensionless or % | Figure 8b geometric washout analysis |
| **Fractional aptamer occupancy** | $\theta$ | 0–1 | dimensionless | Fraction of surface-confined aptamers bound to E2 |
| **Initial fractional occupancy** | $\theta_0$; theta0 | 0.0 | dimensionless | Initial unbound sensing state |
| **Apparent association rate constant** | k_on,app; kon_app_M_inv_s | $5.0 \times 10^5$ | $M^{-1}\ s^{-1}$ | Association term in the occupancy equation; representative simulation input |
| **Default apparent dissociation constant** | K_D,app; Kd_M | 5 nM | M | Default affinity used for Figure 8a,c simulations |
| **Representative apparent dissociation constants** | K_D,app; KD_CASES | 20, 5, 1 | nM | Affinity-dependent heatmaps in Figure 8d–f |
| **Apparent dissociation rate constant** | k_off,app | k_on,app × K_D,app | $s^{-1}$ | Dissociation term during wash; derived from kinetic relation |
| **k_off for K_D = 20 nM** | k_off,app | $1.0 \times 10^{-2}$ | $s^{-1}$ | Lower-affinity heatmap |
| **k_off for K_D = 5 nM** | k_off,app | $2.5 \times 10^{-3}$ | $s^{-1}$ | Default and intermediate-affinity heatmap |
| **k_off for K_D = 1 nM** | k_off,app | $5.0 \times 10^{-4}$ | $s^{-1}$ | Higher-affinity heatmap |
| **Dissociation half-life** | $t_{1/2}$,off | 1.16, 4.62, 23.10 | min | Interprets affinity-dependent regeneration difficulty |
| **Active aptamer surface density** | Γ_T; Gamma_pmol_cm2 | 1.0 | pmol $cm^{-2}$ | Surface capacity and surface uptake/release term; representative input |
| **Total active surface capacity** | A_WE Γ_T | 125.7 | fmol | Total surface binding-site capacity for the modeled electrode |
| **Equivalent site concentration** | A_WE Γ_T / V_ch | 3.29 | nM | Surface capacity expressed relative to chamber volume |
| **Interfacial E2 concentration** | C_s(t) | Simulated algebraic variable | M | Local E2 concentration controlling binding/rebinding at aptamer layer |
| **Reference mass-transfer coefficient** | k_m,ref; km_ref_um_s | 20.0 | μm $s^{-1}$ | Links bulk chamber concentration to interfacial concentration; representative input |
| **Reference flow for k_m scaling** | Q_ref; Q_ref_uL_min | 2.0 | μL $min^{-1}$ | Reference point for flow-dependent mass-transfer scaling |
| **Flow-scaling exponent for k_m** | fixed exponent | 1/3 | dimensionless | Approximate laminar-flow mass-transfer scaling |

**Table 3. Current mapping, numerical settings, and regeneration-output metrics.**

| Parameter | Symbol / code name | Value used | Unit | Use in model |
|---|---|---|---|---|
| **Unbound-state current** | i_U; I_unbound_nA | −230 | nA | Baseline current used in simulated SWV traces |
| **Binding-induced current amplitude** | Δi; DeltaI_nA | 80 | nA | Maps fractional occupancy to current change |
| **Drift term** | drift_nA_per_min | 0.0 | nA $min^{-1}$ | Set to zero to isolate flow-regeneration effects |
| **SWV peak current** | i_p(t) | Simulated output | nA | Figure 8a cyclic current response and Figure 8c metrics |
| **Numerical time step** | dt_s | 5.0 | s | Time step for numerical integration |
| **Integration method** | RK4 | Fourth-order Runge–Kutta | — | Solves coupled chamber and occupancy differential equations |
| **Memory fraction** | residual/response | Derived | dimensionless | Residual current after wash normalized to previous pulse response |
| **Regeneration efficiency** | η_regen | Derived | dimensionless or % | Figure 8c and Figure 8d–f recovery metric |
| **Minimum regeneration efficiency** | min(η_regen) | Derived | % | Conservative heatmap metric across full E2 sequence |
| **Mean regeneration efficiency** | mean(η_regen) | Derived | % | Supplementary design-space metric exported in CSV |
| **Final regeneration efficiency** | η_regen at final dose | Derived | % | Supplementary metric at the 100 nM cycle |
| **Maximum memory fraction** | max(memory) | Derived | % | Worst-case carryover metric exported in CSV |
